# The sets of kinesins and dynein transporting endocytic cargoes determine the effect of tau on their motility

**DOI:** 10.1101/2022.06.10.495679

**Authors:** Daniel Beaudet, Christopher L. Berger, Adam G. Hendricks

## Abstract

The misregulation of tau, a neuronal microtubule-associated protein, is linked to defective axonal transport and neurodegenerative disease. We reconstituted the motility of isolated phagosomes along microtubules to ask how the sets of motors transporting a cargo determine its motility and response to tau. Using quantitative photobleaching, we find that early phagosomes (EPs) and late phagosomes (LPs) are associated with different sets of kinesin-1, -2, -3, and dynein. While EPs exhibit unidirectional retrograde transport, LPs move bidirectionally. Previously, we found that tau biases LP transport towards the microtubule minus-end. Here, we find that tau strongly inhibits long-range retrograde EP motility. Tau impedes the forces generated by multi-dynein teams and accelerates dynein unbinding under load. Thus, specific cargoes differentially respond to tau, where dynein-complexes on EPs are more sensitive to tau than those on LPs.

## Introduction

Motor-mediated transport is essential to maintain the proper localization and function of organelles and vesicular cargoes in cells. Kinesins -1, -2, and -3 are the principal plus-end directed microtubule-based motors that drive anterograde transport, while cytoplasmic dynein drives minus-end directed retrograde transport (Hirokawa et al., 2010; Reck-Peterson et al., 2018). Cargoes bound simultaneously by kinesin and dynein move bidirectionally along microtubules, which enables them to navigate around obstacles and achieve long-range targeted trafficking throughout the crowded cellular environment (Gross et al., 2002; Welte, 2004; Hendricks et al., 2010; Chaudhary et al., 2018). Intracellular cargoes exhibit diverse motility due to differences in their sets of kinesins and dynein motors and the regulatory mechanisms that coordinate their opposing activity. For instance, early endosomes are bound by kinesin-1, kinesin-3, and dynein and typically exhibit short bursts of unidirectional motility (Brown et al., 2005; Hoepfner et al., 2005; Loubéry et al., 2008; Flores-Rodriguez et al., 2011). In filamentous fungi, early endosomes are driven by tightly bound kinesin-3 motors, and the transient binding of dynein causes their directionality to switch between anterograde and retrograde transport (Schuster et al., 2011). Whereas late endosomes and lysosomes that simultaneously associate with members of the kinesin-1, -2, -3 families and dynein exhibit robust bidirectional transport (Matsushita et al., 2004; Brown et al., 2005; Loubéry et al., 2008; Rosa-Ferreira and Munro, 2011; Pu et al., 2016). Multiple types of kinesins on the same cargo could be required for different organelle functions or target organelles to different destinations by controlling the activity of specific kinesins or through their selective transport along microtubules with different posttranslational modifications (Guardia et al., 2016; Pu et al., 2016). Although many kinesin-cargo interactions have been identified (Jenkins et al., 2012; Bentley et al., 2015; Hummel and Hoogenraad, 2021), determining the full complement of native motors on most endogenous cargoes, and how these sets of motors are regulated to achieve targeted trafficking remains a challenging problem.

Microtubule tracks spatially regulate trafficking in cells. Microtubule associated proteins (MAPs) organize the cytoskeleton by controlling the polarity, bundling, and stability of microtubules, as well as the interactions and movement of motor proteins along them (Liu et al., 2012; Lipka et al., 2016; Gumy et al., 2017; Balabanian et al., 2017; Chaudhary et al., 2018; Harterink et al., 2018; Chaudhary et al., 2019; Hooikaas et al., 2019; Monroy et al., 2020; Ferro et al., 2022). Several MAPs regulate the movement of motor proteins. One of the most well-studied is the neuronal MAP tau, which is implicated in Alzheimer’s Disease and related neurodegenerative diseases collectively known as tauopathies. Pathogenic forms of tau perturb the organization of the axonal cytoskeleton and lead to synaptic disfunction and neuronal dystrophy (Mandelkow et al., 2003). Despite the link between aberrant tau regulation and neuropathology, it remains unclear what role tau plays in regulating axonal transport in healthy neurons and how defects lead to neurodegeneration.

Single-molecule studies show that microtubule motors have varying sensitivities to tau. Tau strongly reduces the processivity and landing rates of kinesin-1 and kinesin-3 along microtubules in vitro (Vershinin et al., 2007; Dixit et al., 2008; McVicker et al., 2011; Monroy et al., 2020). In contrast, kinesin-2 is less processive, but is better able to navigate around tau due to its longer more flexible neck linker domain (Hoeprich et al., 2014; Hoeprich et al., 2017). Dynein is also less sensitive than kinesin-1 and kinesin-3 as it often pauses then proceeds to pass-through tau patches along microtubules (Dixit et al., 2008; Vershinin et al., 2008; Tan et al., 2019). In addition, there are multiple isoforms of tau expressed throughout the adult brain that have different impacts on the motility of motor proteins. The 3RS-tau isoform strongly inhibits kinesin-1 and dynein processivity compared to 4RL-tau (Vershinin et al., 2007; Dixit et al., 2008; Vershinin et al., 2008; McVicker et al., 2011). This is likely because the 3RS-tau isoform binds more statically along microtubules and has a higher propensity to form patches compared to 4RL-tau (Hinrichs et al., 2012; McVicker et al., 2014). Tau patch formation is a potential regulatory mechanism to selectively govern the accessibility of the microtubule surface for motors and other MAPs (Tan et al., 2019; Siahaan et al., 2019). Conversely, patches or larger-order tau-complexes that form along microtubules could also act as precursors to tau-aggregates and neurofibrillary tangles found in neurodegenerative disease (Gyparaki et al., 2021). Taken together, these findings suggest that tau functions as a selective barrier along the microtubule lattice that allows some types of motors to pass through while impeding others. Although these studies provide detailed biophysical characterizations of tau’s impact on individual motors, endogenous cargoes are often transported by teams of multiple kinesins and dyneins, and it is less clear how these heterogenous teams of motors are regulated by tau.

MAPs decorate the microtubules that serve as tracks for many types of cargoes, leading us to ask how MAPs might target specific cargoes to different destinations in the cell. To address this question, we characterized the sets of motors bound to endocytic cargoes and used *in vitro* reconstitution to test the impact of 3RS-tau on their motility along microtubules. We made a quantitative comparison of the sets of motors on different cargoes and found that differences in the number and types of motors govern their motility and their regulation by MAPs. In cells, early phagosomes (EPs) are unidirectional and more often transported towards the microtubule minus-end, whereas late phagosomes (LPs) move bidirectionally. We found that the number of kinesin-1 and kinesin-3 motors on EPs and LPs was similar but EPs had fewer dynein and kinesin-2 compared to LPs. In addition, EPs were more heterogenous and bound by fewer motors compared to LPs. Our previous work showed that isolated LPs move bidirectionally *in vitro* and tau biases their transport towards the minus-end of microtubules (Chaudhary et al., 2018). Here we show that tau more strongly inhibits EP motility. Tau reduces the number of long-range minus-end directed transport events but has less of an inhibitory effect on short minus-end or plus-end directed transport. In addition, we show that tau reduces the overall magnitude of forces exerted by EPs with the highest reduction of forces generated by teams of multiple dynein motors. These results suggest that differences in the number and type of motors present on cargoes determine their movement and response to tau. Thus, tau directs the transport of different cargoes by selectively tuning the motility and forces exerted by kinesin and dynein teams.

## Results

### EPs and LPs move differently and are transported by different sets of motors

We first sought to determine how the sets of motors that associate with different cargoes correlate with their motility. To test this, we analyzed phagosomes at different stages of maturation. Latex bead-containing phagosomes are a powerful system to examine how the membrane composition, biochemical properties, and ensembles of motors transporting endocytic organelles change in response to maturation and fusion events (Desjardins et al., 1994a; Desjardins et al., 1994b; Garin et al., 2001; Goyette et al., 2012; Blocker et al., 1997; Loubéry et al., 2008; Rai et al., 2016). Here, we characterized the motility and quantified the sets of motors on early phagosomes (EPs) and late phagosomes (LPs) from mouse J774A.1 macrophage. To image phagosome motility, cells were treated with 200 nm fluorescent latex beads and the movement of bead-containing phagosomes was monitored using total internal reflection fluorescence (TIRF) microscopy. Time lapse images of EPs were recorded 30–45 minutes after bead uptake and LPs were recorded after 90 minutes, which are consistent with time points previously shown to coincide with the biochemical markers of early and late stages of phagosome maturation (Desjardins et al., 1994a; Desjardins et al., 1994b; Garin et al., 2001; Goyette et al., 2012). First, we compared the trajectories of EPs and LPs using TrackMate (Tinevez et al., 2017) (Fig 1A) and plotted their displacement to show directionality towards the cell center (microtubule minus-end) or towards the cell periphery (microtubule plus-end) (Fig 1B). EPs had a higher fraction of long-range, minus-end directed trajectories compared to plus-end directed trajectories, while LPs had an equal fraction of plus-end and minus-end directed trajectories (Fig 1B and C). Phagosomes display stationary, diffusive, and processive periods of motility (Hendricks et al., 2012; Rai et al., 2016). Periods of processive motility were identified as segments of a trajectory between two reversal events with displacements greater than 150 nm, which is consistent with mean-squared displacement (MSD) analysis (Chaudhary et al., 2018). Results show that EPs had a higher fraction of long, minus-end directed processive runs and had fewer reversal events compared to LPs (Fig S1A–C). While most directed trafficking is thought to be driven along microtubule tracks by motor proteins, cargoes could also ‘hitchhike’ along dynamic microtubules to move around in cells. However, dynamic microtubules were previously shown to transport peripheral phagosomes at speeds below 0.1 μm/s compared to motor-driven events that transport phagosomes at faster speeds between 0.2–1.5 μm/s (Blocker et al., 1998), which is consistent with the majority of the processive motility events observed here (Fig S1D). Thus, our results suggest that phagosomes first move towards the perinuclear region of the cell in unidirectional runs, then transition to more bidirectional motility as they progress through maturation to potentially fuse with other organelles and adopt specialized functions (Desjardins et al., 1997; Goyette et al., 2012; Garin et al., 2001).

**Figure 1.**
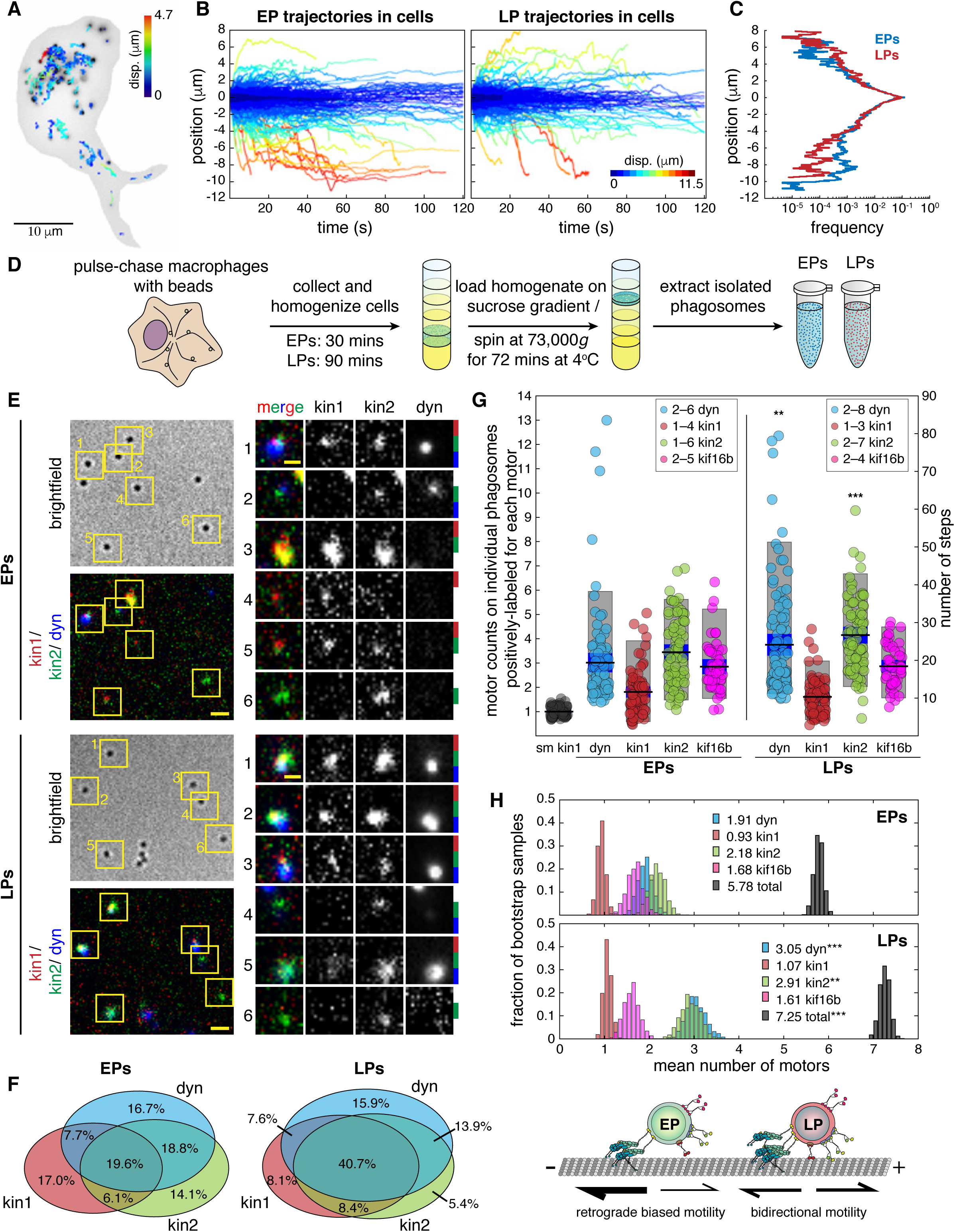
EPs and LPs exhibit diverse motility and are transported by different sets of motors. **A)** An image of a J774A.1 mouse macrophage treated with 200 nm latex fluorescent beads. The image is overlayed with positional tracking data obtained from TrackMate and color coded by displacement. Time-lapse images were acquired 30–45 mins or > 90 mins after bead uptake to visualize bead-containing early phagosomes (EPs) or late phagosomes (LPs), respectively. **B)** The plots show the displacement of EPs (2211 trajectories from 9 cells) and LPs (1579 trajectories from 7 cells) towards the cell center (minus-end direction) or the cell periphery (plus-end direction). **C)** A plot shows the frequency of plus-end and minus-end EP and LP trajectories as a function of displacement. **D)** A schematic shows the steps to isolate phagosomes from cells. EPs were collected at 30 mins and LPs were collected at 90 mins after bead uptake. **E)** Images show isolated EPs (top panel) and LPs (bottom panel) immunolabeled for kinesin-1 (red), kinesin-2 (green), and dynein (blue). On the right, zoomed in ROIs show examples of the different combinations of motors on individual phagosomes. Colored bars on the right indicate the motors present for each example. Scale bars are 2 μm or 1 μm for selected ROIs. **F)** Venn diagrams show the mean percentages of cargo bound by single motors or various combinations of different motors on EPs (n = 230) and LPs (n = 239). **G)** Stepwise photobleaching analysis (Fig. S2) was used to estimate the number of motors that associate with EPs and LPs. A plot shows the distributions of the number of kinesin-1, kinesin-2, kif16b, and dynein motors on EPs and LPs, mean values (black lines), SEM (blue bars), and 90% confidence intervals (light grey bars). Single molecule kinesin-1 (sm kin1) was used to estimate the number of steps for a single motor (n = 59). The step counts indicate that there were 1–4 kinesin-1 on EPs (n = 79) and 1–3 kinesin-1 on LPs (n = 82), 1–6 kinesin-2 on EPs (n = 82) and 1–7 kinesin-2 on LPs (n = 83), 2–5 kinesin-3 on EPs (n = 51) and 2–4 kinesin-3 on LPs (n = 49), and 2–6 dynein on EPs (n = 101) and 2–8 dynein on LPs (n = 107). Bootstrapping of the mean number of steps for each motor was performed to test for statistical significance between EPs and LPs. **H)** Histograms show the mean number of each motor and predicted total number of motors on individual EPs and LPs (See methods). Bootstrapping was used to test for statistical significance and to determine the means and 95% confidence intervals. The schematic depicts the correlation between the sets of motors and the motility of cargo. (** *p* < 0.01, *** p < 0.0001).

The type and number of motor proteins that associate with cargoes governs their transport behavior. Previous studies showed that LPs are transported by teams of kinesin-1, kinesin-2, and dynein motors (Blocker et al., 1997; Hendricks et al., 2012; Chaudhary et al., 2018). However, it was not clear how the sets of motors bound to phagosomes change in response to maturation. To test this, we quantitatively compared the number of kinesins -1, -2, -3, and dynein on EPs and LPs. First, we isolated EPs and LPs from cells and performed 3-color immunofluorescence imaging to detect the combination of motors on individual phagosomes. EPs and LPs were isolated from macrophages at 30 or 90 minutes after bead uptake and purified on a sucrose density gradient (Fig 1D). The largest fraction of LPs (40.7%) were positive for kinesin-1, kinesin-2, and dynein (Fig 1E–F), in agreement with previous results (Chaudhary et al., 2018). In contrast, the motor sets associated with EPs were more diverse and a higher fraction of EPs were found to have only one type of motor compared to LPs (Fig 1E–F). We also probed for kinesin-3 motors on phagosomes. We tested for kinesin-3 motors separately due limitations in the number of fluorophores that can be used to detect multiple motors within the same experiment. EPs and LPs were co-immunolabeled for dynein and the kinesin-3 motors kif16b and kif1b (Fig S1E–F), which have been found to transport similar endocytic organelles (Matsushita et al., 2004; Hoepfner et al., 2005; Blatner et al., 2007). We found that a higher fraction of EPs were positive for kif16b than LPs. The detection levels for kif1b were comparable to non-specific binding controls (Fig S1G–H).

Using stepwise photobleaching analysis, we estimated the number of kinesins -1, -2, -3, and dynein motors on EPs and LPs. Isolated phagosomes were immunolabeled for each motor with Alexa647 antibody separately and imaged using epifluorescence illumination until the fluorescence signal was completely photobleached (Fig S2A). A step-finding algorithm was applied to each intensity trace and used to measure the size and number of steps (Fig S2B–F) (Chen et al., 2014; Chaudhary et al., 2018). Single recombinant kinesin-1 motors were also immunolabeled with Alexa647 antibodies and imaged under similar conditions to determine the number of steps for a single molecule (Fig S2A–C). For those phagosomes that were positively labeled, EPs and LPs had similar numbers of kinesin-1 and kinesin-3 but LPs were more frequently bound by more dynein and kinesin-2 motors (Fig 1G), which is consistent with previous findings (Hendricks et al., 2010; Chaudhary et al., 2018; Cella Zanacchi et al., 2019). To illustrate how a group of motors on a cargo changes as it responds to maturation, we estimated the number of kinesins-1,-2,-3, and dynein motors bound to individual EPs and LPs, taking into account the fraction of phagosomes where a given motor was not detected (see methods). Our predictions indicate that ∼ 50% of LPs were bound by 3 or more different motors, whereas EPs were more often bound by fewer types of motors (Fig S1I). Further, the average individual EP is bound by fewer dynein and kinesin-2 compared to LPs but bound by similar numbers of kinesin-1 and kinesin-3 motors (Fig 1H, S2G–H). Thus, small changes in the number of motors bound to a cargo and therefore the number of motors that can engage along the microtubule can result in large changes in a cargo’s motility (Fig 1H) (Müller et al., 2008; Ohashi et al., 2019).

### Tau inhibits minus-end directed early phagosome motility

We next asked how tau regulates the motility of EPs, which are transported by different sets of motors and move differently than LPs. We previously showed that LPs move bidirectionally along microtubules *in vitro* and that tau biases motility towards the microtubule minus-end (Chaudhary et al., 2018). Here, we tested the impact of tau on the motility of EPs under similar conditions. We reconstituted the motility of isolated EPs along taxol-stabilized, fluorescently labeled microtubules polymerized from bright GMPCPP seeds with or without the addition of 10 nM 3RS-tau. Time lapse recordings of EP motility events were imaged using TIRF microscopy and analyzed by subpixel tracking using 2D-Gaussian-fitting (FIESTA) (Ruhnow et al., 2011). As observed in cells, isolated EPs primarily exhibit unidirectional transport in both directions but had a higher fraction of net motion directed toward the minus-end of microtubules (Fig. 2A–C). With tau, the frequency of long-range minus-end directed EP trajectories was reduced compared to shorter trajectories (< 1 μm) or plus-end directed trajectories (Fig 2A–C). We also noted that the frequency of observable motility events with tau decreased approximately 2-fold, which is likely due to competition for available binding sites between motors and tau (Kellogg et al., 2018; Ferro et al., 2019). In support, linescans of tau and EP max projections show that EPs often bound and proceeded to move in areas of the microtubule where tau levels were low, which indicates that tau reduces the access of EPs to the microtubule surface (Fig 2D). We also examined how tau impacts the motility of those EPs that bind and move along microtubules and found that 44% of plus-end and 67% of minus-end directed EPs paused when they encountered higher levels of tau, whereas 37% of plus-end and 33% of minus-end directed cargoes passed through tau (Fig 2F). Furthermore, 19% of plus-end directed cargoes detached from microtubules when they encountered tau compared to minus-end directed cargoes that remained attached (Fig 2E). These results are in agreement with recent studies that showed that dynein-dynactin-BicD2N complexes are less sensitive to tau and often pass through or pause at tau patches compared to diffusive dynein complexes or kinesin-1 motors that are incapable of passing through tau and detach from the microtubule (Tan et al., 2019; Siahaan et al., 2019; Dixit et al., 2008). Our findings further support a model in which tau acts as a selective barrier to control accessibility of the microtubule surface to different cargoes, as well target their directional transport depending on the sets of motors engaged along them.

**Figure 2.**
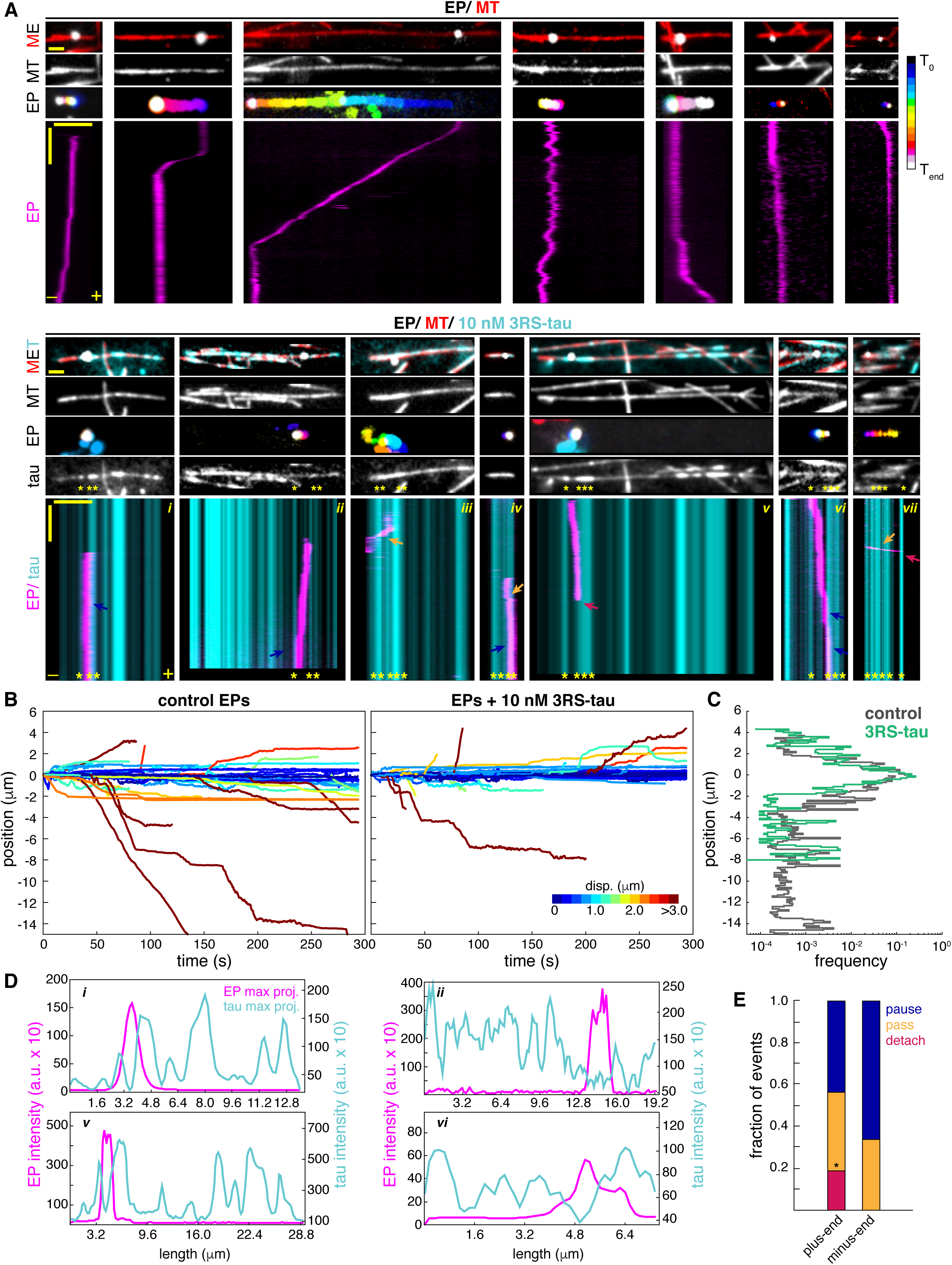
Tau reduces minus-end directed EP motility in vitro. **A)** Early phagosomes were extracted from cells and their motility was reconstituted *in vitro* along polarity marked microtubules (bright segment; minus-end) +/- 10 nM 3RS-tau. Images show microtubules (MT), tau, and max projections of EPs temporally color coded. Kymographs show examples of unidirectional plus-end and minus-end transport, and rare bidirectional transport +/- tau. EPs often pause (blue arrows), and less frequently pass through (orange arrows) or detach from the microtubule (magenta arrows) when they encounter tau patches (yellow asterisks). Horizontal scale bars are 2 μm for images and 5 μm for kymographs, and the vertical scale bars for kymographs are 30 seconds. **B)** Plots show trajectories of EPs directed towards the plus-end or minus-end of microtubules +/- tau (control, n = 55; tau, n = 49). Trajectories are color coded by displacement. **C)** A plot shows the frequency of plus-end and minus-end EP trajectories +/- tau as a function of displacement. **D)** Plots show linescans of the maximum projections of EP and tau fluorescence signals along microtubules. Example plots of images *i, ii, v,* and *vi* from panel A show that EPs often move in areas of the microtubule were tau intensity is low. EP fluorescence intensity is shown on the left y-axis and tau fluorescence intensity is shown on the right y-axis. **E)** A bar graph shows the percentage of EPs that detach, pause, or pass-through tau patches (plus-end events; n = 16, minus-end events; n = 27). The data were analyzed using the Fisher’s exact test (* p < 0.05).

Isolated EPs exhibit periods of stationary, diffusive, and processive transport along microtubules in vitro. To determine the contribution of each mode of transport to their overall motility, we segmented trajectories into periods of stationary, diffusive, and processive transport using change point analysis (Fig 3A). Cargo trajectories were projected along the length of microtubules to track their on-axis position and directionality (Fig 3A). The tracking uncertainty was determined to be 16 nm from the tracked positions of non-motile phagosomes (Fig S3A). Based on the MSD and tracking uncertainty, runs were considered stationary if the run length (R_L_) ≤ 16 nm, diffusive if R_L_ > 16 nm and α ≤ 1, and processive if R_L_ > 16 nm and α > 1 (Fig. 3A and S3B). The MSD for all runs with or without tau indicates that most EP motility is diffusive (control EPs α = 1.0 and EPs with tau α = 0.91) (Fig 3B). The group of runs that were identified as processive by change point analysis had α-values that reflect motor-mediated transport (control EPs α = 1.6 and EPs with tau α = 1.6), whereas runs identified as diffusive had lower α- values (control EPs α = 0.59 and EPs with tau α = 0.61), indicative of constrained diffusion along the microtubule lattice (Fig. 3B). Tau did not change the average lengths of stationary, diffusive, and plus-end directed processive runs but reduced minus-end directed processive runs (Fig 3C), whereas the average velocities for each category remained unchanged (Fig 3D). Moreover, tau did not impact the frequency of stationary and diffusive motility but reduced the frequency of processive transport and altered its net directionality (Fig 3E–F). While stationary and diffusive runs were found to be directionally unbiased, 63% of processive transport was directed towards the minus-end of microtubules in the absence of tau compared to 49% when tau was present (Fig 3F). Combined, these results show that EPs are directionally biased towards the minus-end of microtubules, and tau strongly inhibits minus-end directed motility compared to its effects on plus-end directed motility (Fig 3G).

**Figure 3.**
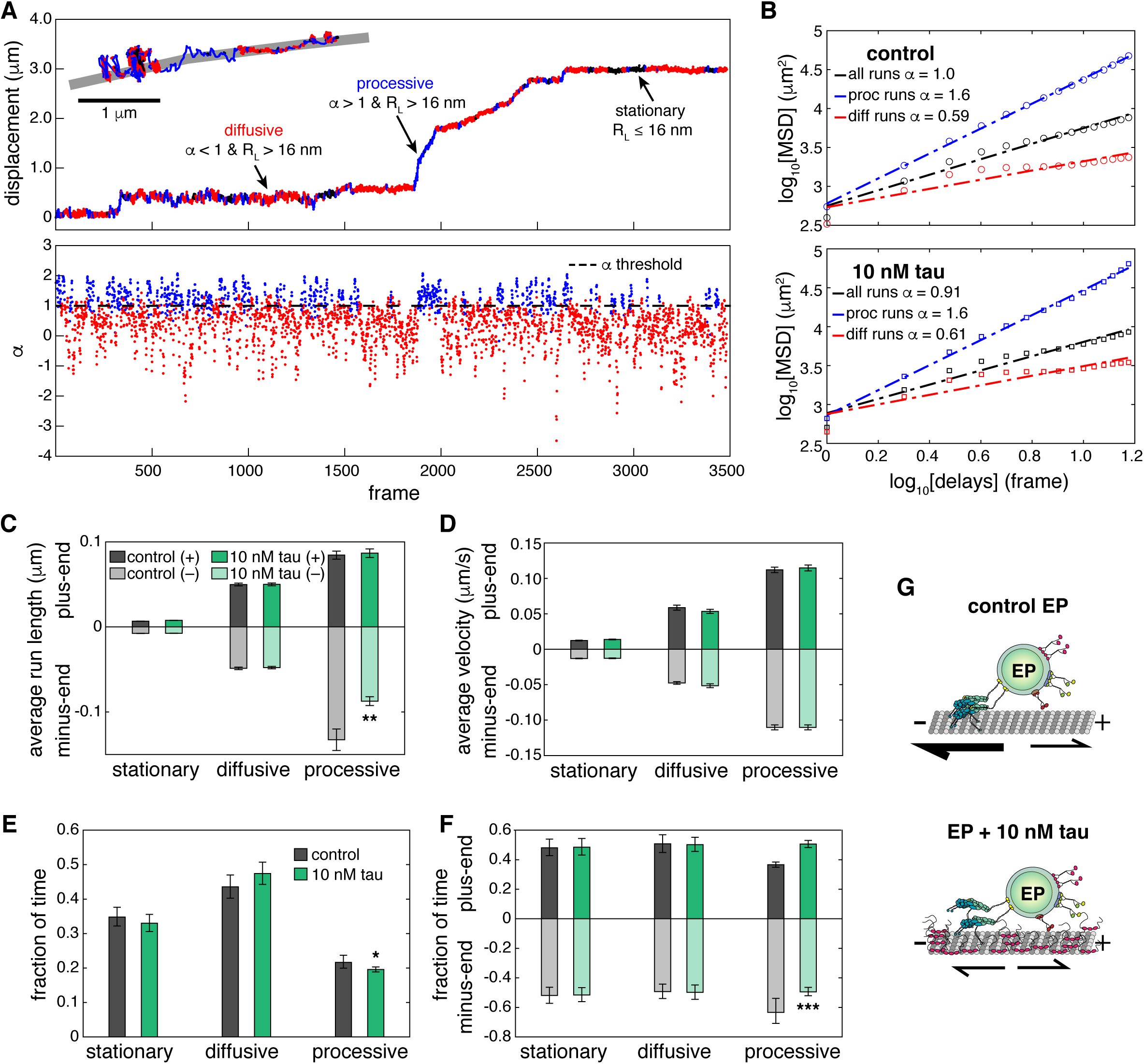
Change point analysis reveals that tau reduces EP minus-end directed processive transport. **A)** Plots show how change point analysis identified stationary (black), diffusive (red), and processive (blue) periods of motility for each trajectory (top graph) by calculating local alpha (α) values using a rolling MSD with a sliding window (bottom graph). The α-value of successive segments must be > 0.3 to be considered a change point. Stationary runs were identified as the period of motility between two change points with a run length (R_L_) ≤ 16nm, which is the calculated mean tracking error (Fig. S3A), diffusive runs were categorized as runs with R_L_ > 16nm and α <= 1, and processive runs were categorized as runs with R_L_ > 16nm and α > 1. The inset shows positional data from tracking an EP along a microtubule using FIESTA. The trajectory is colored to show periods of stationary, diffusive, and processive motility. **B)** The graphs show log-log MSD of EP transport following change point analysis. The α-values are shown for all runs, diffusive runs, and processive runs for control EPs (top) and EPs + tau (bottom). Bar graphs show **C)** the average run lengths and **D)** average velocities for stationary (control (+) n= 863, (-) n= 895; tau (+) n= 791, (-) n= 856), diffusive (control (+) n= 629, (-) n= 616; tau (+) n=680, (-) n= 660), and processive (control (+) n= 748, (-) n= 923; tau (+) n= 885, (-) n= 831) runs identified by change point analysis +/- tau. Error bars show SEM. **E)** A bar graph shows that tau does not significantly change the fraction of time of stationary and diffusive motility but slightly decreases processive motility. **F)** A bar graph shows the fraction of time of stationary, diffusive, and processive runs in the minus-end or plus-end direction +/- tau. Error bars in E and F show 95% confidence intervals. **G)** A cartoon schematic describes the impact of tau on EP motility. Control EPs are unidirectional and biased towards the microtubule minus-end (top). Tau reduces the run lengths and frequency of minus-end directed runs (bottom), which causes EP motility to shift to shorter runs with an equal fraction of plus-end and minus-end directed events. (* *p* < 0.05, ** *p* < 0.001, *** *p* < 0.0001).

### Tau strongly inhibits long-range minus-end directed processive runs

To dissect tau’s role in regulating motor-mediated EP motility, we analyzed the run lengths and velocities of processive runs. Based on the MSD we focused on processive runs with R_L_ > 150 nm (Fig S3B). In addition, tau did not have any noticeable effects on the velocities or run length distributions of stationary, diffusive, and processive runs with R_L_ < 150 nm (Figs S4A–C). The distributions of plus-end and minus-end directed processive runs with R_L_ > 150 nm parsed by length closely follow an exponential decay where long runs were rarer than short runs but more frequently directed towards the microtubule minus-end (Fig 4A). Tau reduced the number of long minus-end directed processive runs, while short runs or plus-end directed runs were unaffected (Fig 4A). Minus-end directed processive runs with lower velocities were also less frequent with tau, whereas plus-end directed velocities did not change (Fig 4B). With tau, the average run length of minus-end directed runs decreased from 0.49 μm to 0.30 μm and the average velocity increased from 0.25 μm/s to 0.28 μm/s, while the average run length and velocity of plus-end runs were not significantly affected (Fig. 4C–D). Next, we used clustering analysis based on the relationship between velocity and run length to determine tau’s impact on plus-end and minus-end directed processive motility (Fig 4E). Plus-end directed runs fit within two distinct groups that proportionally increased in velocity and run length, whereas three groups of minus-end directed runs were identified where the first and second group increase proportionally by velocity and run length, but a third group consisted of runs with longer run lengths and slower velocities. (Fig 4E). Tau significantly reduced the number of runs within the group of slow, long-range minus-end directed runs compared to its effects on the groups of faster, shorter minus-end runs or plus-end directed runs (Fig 4E). These results suggest that for minus-end directed cargoes like EPs, tau acts as an obstacle on the microtubule that more strongly inhibits long dynein-driven runs compared to the relatively short and infrequent kinesin-driven runs.

**Figure 4.**
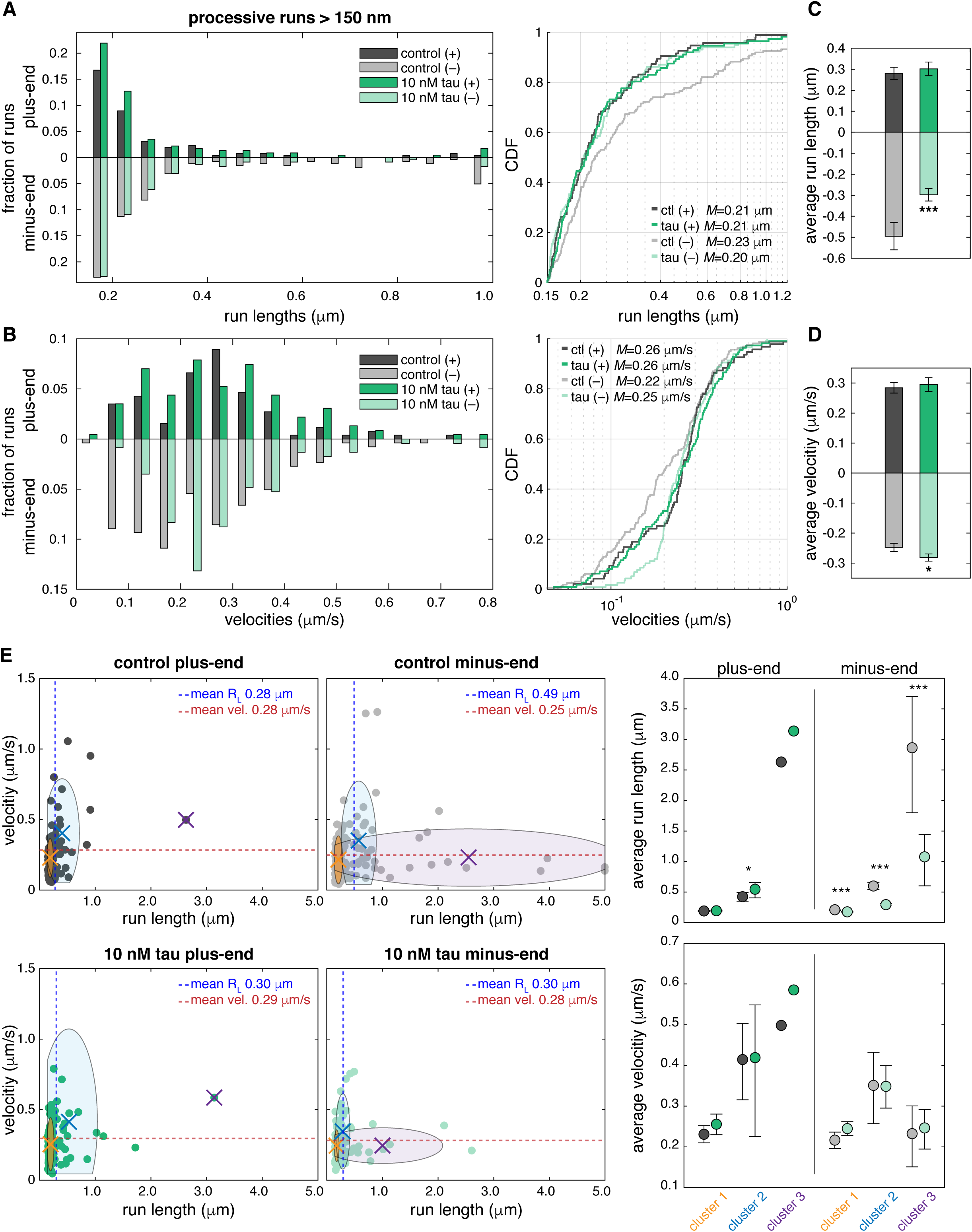
Tau strongly inhibits long-range minus-end directed processive runs. **A)** A histogram shows the distribution of run lengths for processive runs > 150 nm in the plus-end and minus-end direction +/- tau. Analysis of processive runs > 150 nm consisted of 95 plus-end and 162 minus-end control runs and 112 plus-end and 116 minus-end runs with tau. The cumulative distribution function (CDF) plot on the right shows that tau decreases the number of long minus-end directed processive runs but does not impact plus-end directed runs. (plus-end runs, not significant (n.s.); minus-end runs, *p* < 0.05). **B)** A histogram shows the distribution of velocities for processive runs > 150 nm in the plus-end and minus-end direction +/- tau. The CDF plot on the right shows that tau decreases the fraction of runs with slower velocities but has less impact on the fraction of runs with higher velocities or plus-end directed runs (plus-end runs, n.s.; minus-end runs, *p* < 0.0001). Median (*M*) run lengths and velocities are shown in A and B. The Kolmogrov-Smirnov test was used to test for statistical significance of tau’s effects on the distribution of run lengths and velocities. Bar graphs show **C)** the average run lengths and **D)** average velocities for plus-end and minus-end directed processive runs > 150 nm +/- tau. Error bars indicate SEM. **E)** Plots show velocities vs. run lengths of plus-end and minus-end directed processive runs for control (top) and + tau (bottom). Clustering analysis was used to identify groups of runs by velocity and run length. Plus-end directed runs fit within two clusters, whereas minus-end directed runs fit within three clusters. On the right, plots show the average run length and velocity of each cluster. Minus-end directed runs with low velocities and longer run lengths are more strongly inhibited by tau compared to shorter faster runs or plus-end directed runs. (* *p* < 0.05, *** *p* < 0.0001).

### Tau reduces the forces generated by teams of multiple dyneins transporting EPs

To understand how tau regulates the activity of teams of kinesin and dynein, we measured the forces generated by motors transporting EPs along microtubules using an optical trap. Previous studies showed that isolated LPs exert maximum forces of 9–12 pN in vitro and up to ∼20 pN in live cells (Rai et al., 2013; Hendricks et al., 2012; Chaudhary et al., 2018; Chaudhary et al., 2019). The maximum forces generated by the sets of motors on isolated EPs reached stall forces of ∼8 pN directed towards the microtubule minus-end and ∼5 pN directed towards the microtubule plus-end (Fig 5A–B, S5A–B), consistent with fewer motors being associated with EPs than LPs (Fig. 1F–H). EPs rapidly bind to microtubules and exert unidirectional forces directed towards the plus-end or minus-end of microtubules with rare occurrences of phagosomes exerting bidirectional forces (9/63 EPs showed bidirectional forces) (Fig 5A). Consistent with their motility, EPs had a higher fraction of minus-end directed events compared to plus-end directed events (Fig 3F and S5C). Plus-end events occurred over a range of forces with forces above ∼4 pN occurring less frequently than lower force events (Fig 5A and S5A). The higher forces observed are consistent with the stall forces of a single kinesin-1 or kinesin-2 motor (Schroeder et al., 2012), while the higher fraction of lower forces between ∼1–3 pN indicate that kinesins often detach from the microtubule before maximum stall forces are reached (Fig 5B). Alternatively, low-force events might also be driven by kinesin-3 motors (Budaitis et al., 2021). Minus-end forces frequently had lower magnitudes that ranged between ∼1–3 pN, which are consistent with sub-stall detachments or low-force stalls by a single dynein motor (Rai et al. 2013, Chaudhary et al. 2018; Belyy et al., 2016). There were also rare minus-end events with higher forces between ∼5–8 pN (Fig 5B). In the presence of tau, force events directed towards the minus-end of microtubules occurred less frequently and the overall magnitude of forces in both directions decreased (Fig 5C–D, S5C). Tau decreased the frequency of plus-end maximum stall forces above 4 pN, while the frequency of lower force events remained unchanged (Fig 5C–D). However, minus-end directed forces > 5 pN were completely abolished with tau, followed by a moderate increase in the frequency of lower forces at ∼3 pN (Fig 5C–D, S5B).

**Figure 5.**
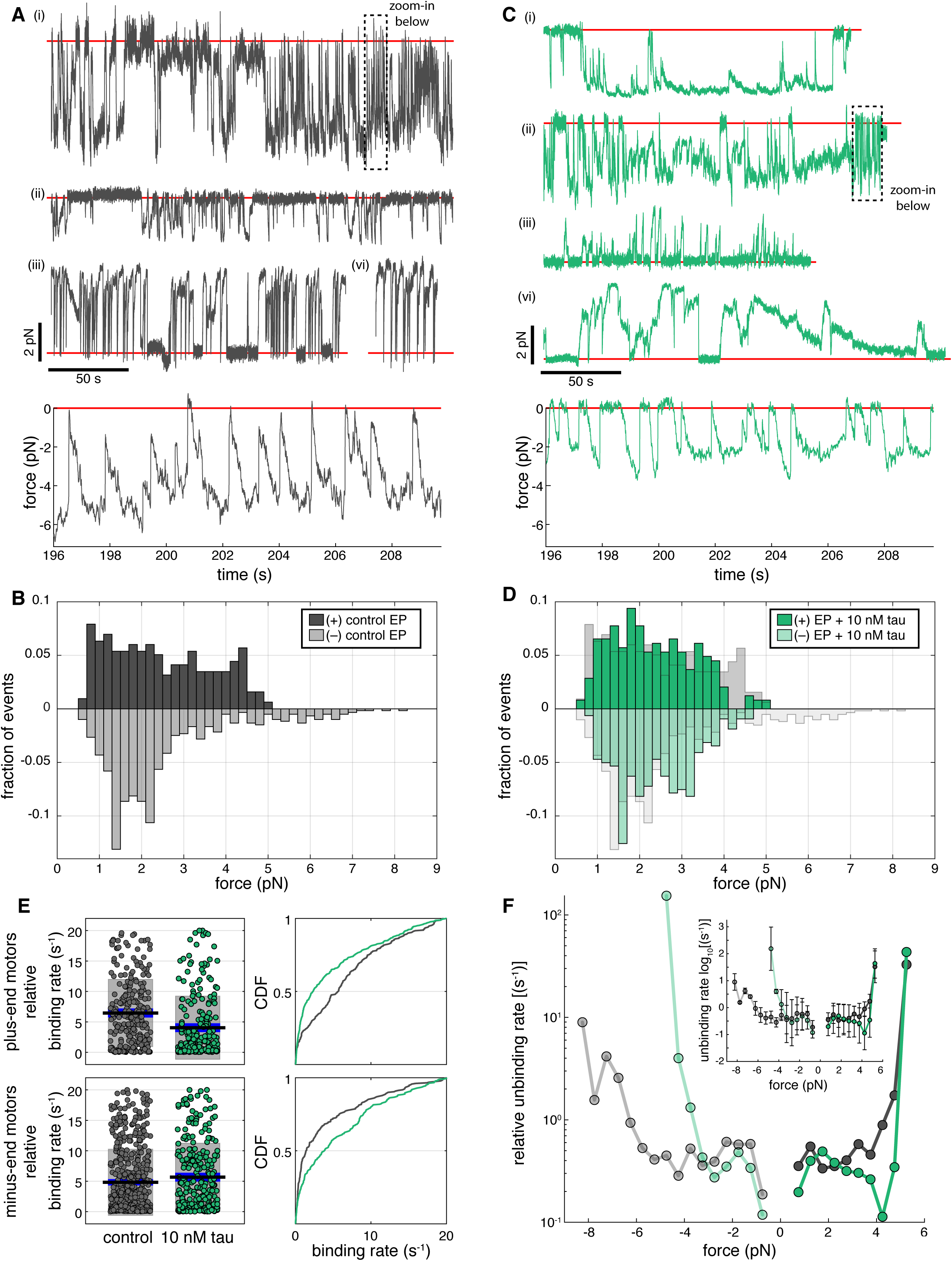
Tau reduces the forces generated by teams of multiple dynein transporting EPs. **A** and **C)** Force traces of EPs +/- 10 nM tau were acquired at 2 kHz using an optical trap and median-filtered at 20 Hz. Force events were observed in both the minus-end direction (*i*, *ii*) and the plus-end direction (*iii*, *iv*) for A) control EPs (n = 904 events from 64 recordings taken over 11 independent experiments) and C) EPs + tau (n = 563 events from 54 recordings over 11 experiments). Red lines show the trap center (0 pN). **B** and **D)** Histograms show the distribution of plus-end and minus-end force events with stall durations > 250 msec for B) control EPs and D) EPs + tau. Control EP force events were overlaid (light grey bars) on top of EP+ tau force events to show how tau reduces higher force events in both directions (plus-end forces, *p* < 0.05; minus-end forces, *p* < 0.001). **E)** Plots show the relative binding rates calculated for plus-end motors (top) and minus-end motors (bottom). On the right, CDF plots show how tau influences the fraction of the relative binding rates of plus-end and minus-end motors (plus-end motors, *p* < 0.0001; minus-end motors, *p* < 0.05). The Kolmogrov-Smirnov test was used to test for statistical significance of tau’s effects on the distribution of forces and relative binding rates. **F)** A plot shows the force-dependent relative unbinding rates of minus-end and plus-end directed forces +/- tau with stall times > 250 msec. The inset shows the log-unbinding rates on the y-axis. error bars indicate SEM.

To compare the effects of tau on the rate at which kinesin and dynein bind to microtubules, we calculated the relative binding rates from the intervals of diffusive dwell times that occurred before each force event in the optical trap (Fig S5D) (Chaudhary et al., 2019). Tau decreased the relative binding rate of kinesin, whereas the binding rate of dynein increased slightly (Fig 5E). In addition, we compared the effects of tau on the rate at which kinesin and dynein detach from microtubules, by using the method described by Berger et al. (2019), which estimates the force-dependent unbinding rates from the probability that the motors detach within a given range of forces, with the simplifying assumption that the force dependence of the unbinding rate is exponential. We found that the unbinding rate for plus-end and minus-end force events with stall durations > 250 msec increased exponentially with force (Fig 5F). However, a higher rate of unbinding also occurred at lower forces (< 1 pN), which indicates that motors transporting EPs frequently unbind from the microtubule independent of force (Fig S5D). Tau increased the unbinding rate of minus-end directed forces, particularly at forces > 3 pN (Fig 5F). In contrast, tau did not affect plus-end forces between 0.5–4.0 pN or significantly alter the unbinding rate (Fig 5E). The lack of an effect of tau on plus-end directed forces is likely due to kinesin-driven forces being rare, consistent with the predominantly minus-end directed motility of EPs (Fig 3). In contrast to bidirectional cargoes like LPs, the dynein teams on minus-end directed unidirectional cargoes are more sensitive to tau on microtubules and often detach before maximum forces are exerted.

## Discussion

Our results show that tau regulates organelle motility in a cargo-specific manner. Early phagosomes exhibit fewer long, minus-end directed runs in the presence of tau while tau enhances dynein-directed movement on late phagosomes. Neuronal cytoskeletons are dense and highly complex, which makes it difficult to dissect molecular mechanisms from observations in cells. Therefore, we used *in vitro* reconstitution and single molecule approaches to understand how tau impacts the transport of endogenous cargoes. This system enabled us to measure the motility of phagosomes and use an optical trap to probe for the biophysical effects of tau on their trafficking along microtubules in the absence of other cellular components. Combined, our data supports a model where tau acts as a selective barrier along microtubules to control the transport of cargoes bound by different sets of motors. We compared early and late phagosomes and found that the types of motors associated with EPs are more variable than LPs, and that EPs associate with fewer dynein and kinesin-2 motors (Fig. 1E-H). The differences in the sets of motors associated with them alters their motility and response to tau (Fig 1B, 2). LPs move bidirectionally (Fig 1B–C), and we showed previously that their plus-end directed forces decreased in the presence of tau, which concomitantly increased the processivity and opposing forces generated by dynein motors along microtubules (Chaudhary et al., 2018). Here, we found that EPs are unidirectional and were more frequently transported towards the minus-end of microtubules (Fig 1B–C, 2A–B). Tau reduced the processivity and maximum forces exerted by teams of dynein on EPs (Fig 4 and 5), which decreased the frequency and travel distances of their minus-end directed transport (Fig 2–4). These findings reveal how tau differentially regulates teams of motors transporting different cargoes (Fig 6). Further, our results suggest that perturbations to tau would be expected to have disparate effects on different types of cargoes.

**Figure 6.**
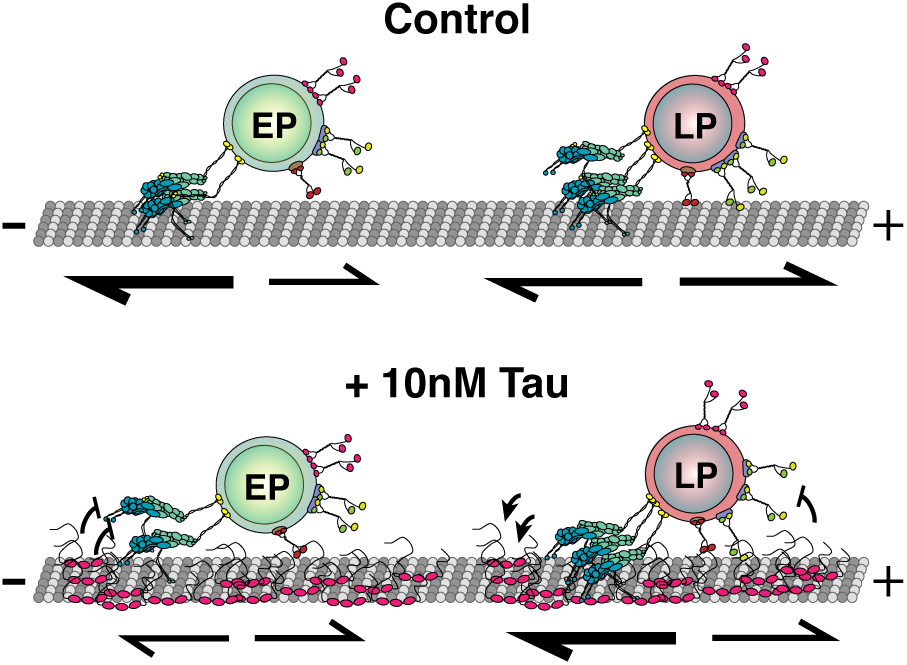
Tau differentially regulates the teams of motors transporting different cargoes. A cartoon schematic shows how EPs and LPs have disparate responses to tau on microtubules. While LPs are bidirectionally transported along microtubules, EPs exhibit unidirectional motility and more frequently move towards the minus-end of microtubules. Tau differentially impacts the motility of EPs and LPs. In the presence of tau, LP transport is biased towards the microtubule minus-end, whereas EP long-range minus-end directed transport is strongly inhibited.

### What determines if a cargo moves unidirectionally or bidirectionally?

Our results are consistent with bidirectional motility resulting from a stochastic tug-of-war between opposing motors that are simultaneously bound to a cargo (Müller et al., 2008). Stochastic tug-of-war models predict that altering the number of kinesin and dynein motors that are bound to a cargo or adjusting their microtubule binding kinetics determines the extent of bidirectional and unidirectional transport (Müller et al., 2008; Soppina et al., 2009; Hendricks et al., 2010; Chaudhary et al., 2018; Ohashi et al., 2019). We reason that EPs exhibit unidirectional motility because they are typically bound by a single type of motor or combinations of two different motors, whereas LPs move bidirectionally because they are simultaneously bound by multiple copies of kinesins-1, -2, -3 and dynein (Fig 1E–F, S2A–B). Alternatively, EPs might be regulated such that opposing motors are not in an active state. In support, our optical trapping data shows that EPs exert unidirectional forces that are comparable to the forces produced by a single kinesin or small teams of dynein motors with rare events driven by larger teams of dynein motors (Fig 5A–B), whereas the magnitude of the forces exerted by LPs were previously shown to be approximately equal in both directions, which corresponds to forces produced by teams of up to 3 kinesin and 10 dynein motors competing in a tug-of-war along microtubules (Chaudhary et al., 2018). These results further indicate that only a subset of the total number of motors that are bound to cargoes are engaged with microtubules and exert force on EPs. Combined, this data suggests that as phagosomes mature, their total number of motors as well as the ratio of kinesin and dynein motors that engage the microtubule change, which potentiates their transition from unidirectional to bidirectional transport.

While our data show that early and late phagosomes exhibit different motility due to being associated with different sets of kinesin and dynein motors, other factors likely modulate the activity of vesicle-bound motors. In *Dictyostelium,* phagosomes examined at 5–10 and 30 minutes following bead uptake were shown to transition from bidirectional to unidirectional retrograde transport (Rai et al., 2016). As phagosomes mature, multiple dynein motors cluster within cholesterol-rich microdomains along their surface, which enables them to collectively exert higher forces (Rai et al., 2016). Thus, clusters of dynein can more effectively outcompete single diffuse kinesins, which promotes the transition from bidirectional to unidirectional retrograde transport. Furthermore, transport may also be differentially controlled in specific regions of the cell by transient motor-cargo interactions. For example, the directionality of early endosomes in filamentous fungi is governed by the transient binding of dynein that outcompetes kinesin-3 motors causing them to switch between plus-end and minus-end directed transport as they approach the hyphal tip, where pools of dynein are most concentrated (Schuster et al., 2011). Similar mechanisms could ensure that phagosome trafficking is spatially and temporally controlled as they mature, where different motors transiently interact with EPs and LPs or are reorganized on their membranes to bias their trafficking in different regions of the cell.

Adaptor proteins are functionally diverse and associate with different motors to control cargo motility. Adaptor-motor complexes have been shown to recognize different cargoes through direct or indirect interactions with Rab GTPases and other membrane associated regulatory effectors (Cantalupo et al., 2001; Jordens et al., 2001; Niwa et al., 2008; Schonteich et al., 2008; Arimura et al., 2009; Horgan et al., 2010; Ueno et al., 2011; Christensen et al., 2021). As phagosomes mature, the level of Rabs and other small GTPases change (Goyette et al., 2009). Therefore, it is possible that these changes generate downstream effects that alter the type of adaptors and motors that associate with early and late phagosomes, which in turn impact their motility. Adaptor proteins orchestrate bidirectional motility or enhance transport by scaffolding multiple kinesin and dynein motors (Akhmanova and Hammer et al., 2010; Fu and Holzbaur, 2013; Bielska et al., 2014; Kendrick et al., 2019; Fenton et al., 2021). Adaptors also modulate the effect of tau on motor motility. TRAK1, which is a mitochondrial specific adaptor protein, is an example of an adaptor that enhances anterograde transport, where it activates and tethers kinesin-1 to the microtubule surface enabling it to robustly transport mitochondria over long distances and through dense tau patches along microtubules (Henrichs et al., 2020). While several adaptors have been identified to control the motility of different organelles and vesicular cargoes, those that associate with phagosomes are not fully characterized. Yet, different adaptor-motor complexes likely associate with EPs and LPs, and contribute to differences in their motility and responses to tau by regulating both the recruitment and activity of phagosome-bound motors.

### How are cargoes with different sets of motors selectively regulated along tau-decorated microtubules?

Our work illustrates that the response of organelle motility to tau is not fully predicted from the response of single motor proteins to tau. Previous studies investigated tau’s impact on the motility of single motors or groups of homogenous motors and showed that kinesin-1 and kinesin-3 are more strongly inhibited by tau than kinesin-2 and dynein (Vershinin et al., 2007; Dixit et al., 2008; Vershinin et al., 2008; McVicker et al., 2011; Hoeprich et al., 2014; Hoeprich et al., 2017; Monroy et al., 2018; Tan et al., 2019). It is becoming more clear that tau acts like a roadblock for certain motors depending on their structure as well as their microtubule binding kinetics. Kinesin-1 predominantly moves by stepping forward toward the plus-end of microtubules along a single protofilament, while dynein moves with less precision but is capable of side-stepping to adjacent protofilaments and taking backwards steps (DeWitt et al., 2012). Kinesin-2 is also capable of side-stepping due to its more flexible neck linker (Shastry and Hancock, 2010; Hoeprich et al., 2014; Hoeprich et al., 2017). Because of their less restrained movement, it is thought that kinesin-2 and dynein are better at navigating through tau patches compared to kinesin-1 and kinesin-3 that move more rigidly along microtubules. Thus, single-molecule studies would predict that dynein-driven transport would be less sensitive to tau compared to kinesin-driven transport, while we observe that the long-range transport of EPs mediated by dynein is strongly inhibited by tau.

Emerging evidence also suggests that motor detachment and reattachment kinetics play a pivotal role during transport and could influence how cargoes respond to MAPs. Kinesin-1 provides more sustained force during transport but reattaches slowly after it detaches from the microtubule, whereas kinesin-2 and kinesin-3 frequently detach from microtubules under load but reattach much faster than kinesin-1 (Feng et al., 2018; Arpağ et al., 2019). Because their binding rates are much faster than their unbinding rates, kinesin-2 and kinesin-3 can more efficiently probe for available landing sites along microtubules, which helps tether the cargo close to the microtubule and navigate around obstacles. These studies could explain how cooperation between motors, for instance those that produce force and others that may act as a tether, enable cargoes to endure long distance travel over several microns (Fig 2A–B). Although EPs travel further distances, because they are bound by fewer types of motors, they may be more prone to inhibition by tau compared to cargoes that are bound by multiple types of motors like LPs, in which more kinesin and dynein motors engage the microtubule simultaneously to bypass tau (Chaudhary et al., 2018).

In neurons, several mechanisms must coordinate to ensure that cargoes are delivered with precision over long distances. MAPs that differentially impact the motility of kinesin and dynein motors compete for space along microtubules to govern the transport of intracellular cargoes (Monroy et al., 2018; Chaudhary et al., 2019; Monroy et al., 2020). Microtubules within the axon that are decorated by tau could modulate directional transport or function as selective tracks for certain cargo, whereas microtubules that are bound by different MAPs act as selective tracks for other types of cargo. Our results suggest that tau functions as a barrier along microtubules to selectively inhibit some cargoes while allowing the passage of others depending on the type of motors and adaptors driving their transport. Tau is also regulated by post-translational modifications and isoform expression, which could influence its affinity for microtubules as well as its impact on the motility of different motors. Our studies indicate that regulating tau dynamics through post-translational modifications or disease mutations would be expected to have cargo-specific effects on transport.

## Author Contributions

DB and AGH conceptualized and designed this work. DB performed experiments and analysis. DB and AGH wrote the manuscript and made the figures, CLB contributed reagents, CLB and AGH obtained funding for this work.

## Star Methods

### Resource availability

#### Lead contact

Further information and requests for resources and reagents should be directed to and will be fulfilled by the lead contact, Adam G. Hendricks.

### Experimental model and subject details

#### Cell culture

J774A.1 mouse macrophage (ATCC) were maintained in DMEM (Gibco), supplemented with 10% fetal bovine serum (Thermo Fisher Scientific) and 1% GlutaMAX (Gibco) at 37°C with 5% CO_2_.

#### Phagosome isolation

Macrophages were plated in 4 10 cm cell culture dishes and grown to ∼80% confluency at 37°C with 5% CO_2_. Phagosome isolation was performed as described previously (Hendricks et al., 2014). Beads were coated with 10% BSA and incubated in cells for 30 mins or 90 mins to isolate EPs or LPs, respectively. While 200 nm fluorescent beads were used for motility assays, larger 500 nm beads (yellow-green fluorescent; F8813, Invitrogen) were used for optical trapping and non-fluorescent beads were used for immunofluorescence imaging experiments. After treatment with beads, cells were washed with cold PBS and collected using a cell scrapper and spun at 650*g* at 4°C for 5 mins. Cells were resuspended in motility assay buffer (MAB; 10 mM PIPES, 50 mM K-Acetate, 4 mM MgCl_2_, and 1 mM EGTA, pH 7.0.) supplemented with protease inhibitor cocktail (PIC; BioShop), 10mM DTT, 1mM MgATP and 8.5% sucrose and transferred to a tight-fitting Dounce cell homogenizer and homogenized on ice by hand with 5– 10 up-and-down strokes to ensure adequate disruption of cell membranes. Following, the cell homogenate was spun at 650*g* and ∼1 ml of supernatant was mixed with 62% sucrose solution at a 1:1.2 ratio. All sucrose solutions were supplemented with PIC, 10 mM DTT, and 1 mM MgATP. The homogenate mixture was loaded on top of a 3.92 ml 62% sucrose cushion in an Ultra-Clear 5/8” X 4” Beckman Coulter tube (344061). The remaining sucrose solutions: 2.61 ml 35% (or 30% for 200 nm beads), 2.61 ml 25%, and 2.61 ml 10% were added on top of the ∼2 ml homogenate mixture. The sucrose gradient was centrifuged in a swinging bucket rotor at 73,000*g* for 72 mins at 4°C. The bead containing phagosomes appeared as a thin band at the 10–25% (or 25–30% for 200nm beads) sucrose interface and extracted using a 21-gauge needle and syringe. Isolated phagosomes were kept on ice at 4°C or flash frozen and stored at - 80°C.

#### Polarity-marked microtubules

Bright double-cycled GMPCPP microtubule seeds were prepared as described in Hyman et al., (1991), by mixing 25% Alexa647 labeled tubulin and 75% unlabeled tubulin in BRB80 (80 mM K-PIPES, 1 mM MgCl_2_, 1 mM EGTA, pH 6.8) to a final concentration of 5 mg/ml supplemented with 1 mM GTP (Sigma Aldrich) and polymerized at 37°C for 20 mins. Bright microtubule seeds were then vortexed in cold BRB80 supplemented with 1 mM GMPCPP (Jena Bioscience), 1 mM MgSO_4_, and 2 mM MgCl_2_ and incubated on ice for 5 mins. This mixture was incubated at 37°C for 30 mins and pelleted at 30 psi for 5 mins in the Beckman Airfuge using the A95 rotor. The pellet was resuspended in 70 μl cold BRB80 and depolymerized on ice for 20 mins. The solution was mixed with 1 mM MgSO_4_, 2 mM MgCl_2_, and 1 mM GMPCPP and incubated on ice for 5 mins. The mixture was polymerized at 37°C for 30 mins then spun at 30 psi for 5 mins. The pellet was resuspended in BRB80 and aliquoted in 5 μl stocks, flash frozen and stored at -80°C. To prepare polarity marked microtubules, GMPCPP seeds were warmed at 37°C for 1 min and added to a mixture of 8% labeled tubulin and 92% unlabeled tubulin to a final concentration of 2 mg/mL in BRB80 and 1 mM GTP and incubated for 25 mins at 37°C. Microtubules were then stabilized with 20 μM Taxol (Cytoskeleton) and incubated for 25 mins at 37°C. Microtubules were cleared 2X by pelleting them at 10,600*g* for 5 mins at RT then washed with T-BRB80 (BRB80 supplemented with 20 μM Taxol).

### Method details

#### Live cell imaging

Macrophages were plated in MatTek 35 mm glass-bottom dishes (no.1) and grown to ∼50% confluency at 37°C with 5% CO_2_. To perform live imaging, cells were treated with 200 nm blue fluorescent latex beads (F8805, Invitrogen) coated with 10% BSA diluted 1:100 in DMEM media and incubated at 37°C with 5% CO_2_. To image early phagosomes (EPs) cells were incubated for 30 mins with beads prior to imaging and to image late phagosomes (LPs) cells were incubated for 90 mins. Following, cells were washed 1X with PBS to remove non-phagocytosed beads and media was replenished with phenol red-free Leibovitz L-15 Media (Gibco). Live cell imaging was performed on an Eclipse Ti-E inverted microscope (Nikon) with custom optics for total internal reflection fluorescence (TIRF) and imaged using EMCCD camera (iXon U897; Andor Technology) maintained at 37°C. Time lapse recordings were acquired with 30 msec exposures using a 450 nm laser (100 mW) set at 2% power for 120 sec per cell using NIS-Elements acquisition software (Nikon). Image files were exported as TIFFs, which were opened with ImageJ (NIH) and bead containing phagosomes were tracked using TrackMate (Tinevez et al., 2017). Phagosomes were first identified using a Laplacian of Gaussian filter to detect the fluorescence signals. Detected spots were then tracked with the Linear Assignment Problem tracker and linked from frame-to-frame to trace trajectories of EPs and LPs. The tracking parameters were set to link tracks with distances < 0.5 μm and with gaps of two frames if the cargoes were temporarily out of frame. The *xy*-coordinates of each trajectory were imported into MATLAB (The MathWorks) to analyze the mode of transport, directionality, displacement, and number of reversals for all trajectories within cells using custom scripts.

#### Immunofluorescence imaging

Isolated early and late phagosomes were added to silanized coverslips mounted to microscope slides using vacuum grease and double-sided tape to make flow chambers (Dixit and Ross, 2010) and incubated for 1 hour at room temperature (RT) to allow them to adhere to the surface. Following the incubation, chambers were washed 2X with MAB before and after treatment with Pluronic F-127 to prevent non-specific binding before incubation with primary antibodies. Phagosomes were incubated with different combinations of primary antibodies for kinesin-1 (MAB1614, EMD Millipore), kinesin-2 (ab24626, Abcam) conjugated to Alexa568 (A20184, Thermo Fisher Scientific), dynein (sc-9115, Santa Cruz), kif16b (SAB1401759, Sigma), and kif1b (MABC309, EMD Millipore) for 1 hour at RT in the dark. Following treatment with primary antibodies, phagosomes were washed 3X with MAB and incubated with secondary antibodies against mouse (Alexa488; A11029, Thermo Fisher Scientific) and rabbit (Alexa647; A32733, Thermo Fisher Scientific) for 1 hour at RT in the dark. Samples were washed 3X with MAB before imaging. Multichannel images were acquired in brightfield or with 500 msec exposures in TIRF using the 488 nm, 561 nm, and 640 nm lasers, set between 1–5% power. Thresholds > 800 a.u. were applied to compare the type of motors on EPs and LPs. Only phagosomes with fluorescence signals above the threshold were counted.

#### Stepwise photobleaching

Phagosomes were incubated in flow chambers for 1 hour and treated with Pluronic F-127 as described above. Separate chambers were used for each motor counted. Phagosomes in each chamber were incubated with one of the following mouse monoclonal antibodies against kinesin-1 (MAB1614, EMD Millipore), kinesin-2 (ab24626, Abcam), kif16b (SAB1401759, Sigma), kif1b (MABC309, EMD Millipore), and dynein (MAB1618, EMD Millipore) for 2 hours at RT in the dark. Chambers were washed 3X with MAB and incubated with anti-mouse Alexa647 (A21236, Thermo Fisher Scientific) secondary antibody for 1.5 hours. Following, chambers were washed 3X with MAB. Phagosomes were imaged in epifluorescence with 500 msec exposures using a 640 nm laser at 3 mW. Recombinant rat kinesin-1 (rkin430-GFP; A gift from Dr. Gary Brouhard, Dept. Biology, McGill University) was imaged under similar conditions to determine the number of steps for a single motor. Purified kinesin-1 was diluted to 10 nM and incubated with primary and secondary antibodies and bound to unlabeled microtubules using 1 mM AMPNP to detect single statically bound molecules. Images were acquired similar to phagosomes but instead using TIRF to achieve detectable steps. Only steps > 15 a.u. were considered. For kinesin-2, the number of steps was multiplied by a factor of 2 since it exists as a heterodimer and contains 1 epitope recognized by the antibody. A step-finding algorithm (Chen et al., 2014) was used to determine the number and size of photobleaching steps for single motors and for the sets of motors on phagosomes. The mean step size was determined to be 21 a.u. for kinesin-1, kinesin-2, kif16b, and dynein, and 52 a.u. for recombinant single molecule kinesin-1. Steps within the last half of each photobleaching trace were better resolved and therefore only considered for estimating step sizes. The number of steps was determined by dividing the difference between the initial and final fluorescence intensity by the mean step size.

#### In vitro motility assays

Flow chambers were first incubated with anti-!3-tubulin (T4026 clone TUB2.1, Sigma) diluted 2:50 in BRB80 for 5 mins. Chambers were then treated with F-127 for 5 mins and washed 2X with T-BRB80. Polarity marked microtubules were added to the chambers and incubated for 5 mins at RT. Unbound microtubules were washed out with 2X T-BRB80. Isolated phagosomes were flown through the chamber supplemented with 0.2 mg/ml BSA, 10 mM DTT, 1 mM MgATP, 20 μM Taxol, 15 mg/ml glucose, ≥ 2000 units/g glucose oxidase, ≥ 6 units/g catalase, and 1 mg/ml casein. For experiments with tau, microtubules were incubated with 10 nM of Alexa568 labeled 3RS-tau for 30 mins. Tau protein was expressed and purified as previously described in Chaudhary et al., (2018). Phagosome mixtures were also supplemented with 10 nM tau. Using TIRF, microtubules and tau were imaged for 1 frame with the 561 nm and 640 nm lasers at 1–10% power and time lapse recordings of phagosomes were imaged with 80 msec exposures using a 450 nm laser set at 2% power for 5 mins.

#### Subpixel tracking and change point analysis

Motility events were analyzed using FIESTA (Fluorescence Image Evaluation Software for Tracking and Analysis) (Ruhnow et al., 2011). Phagosome fluorescence signals were fitted using the symmetrical 2D-Gaussian function and trajectories analysed using custom MATLAB scripts. Change point analysis was applied to identified periods of stationary, diffusive, or processive periods of motility for each trajectory by finding the local α-values using a rolling MSD analysis over a sliding window length of 12 frames. Change points occurred when the difference between two adjacent α-values was > 0.3 with a minimum of 3 frames between them. The tracking uncertainty was determined to be 16 nm from the tracked positions of non-motile phagosomes (Fig S3A). Runs were identified as stationary (R_L_ ≤ 16 nm), diffusive (R_L_ > 16 nm and α ≤ 1), or processive (R_L_ > 16 nm and α > 1). Runs were further categorized as plus-end directed or minus-end directed, and analysed to determine the velocities, run lengths, and time intervals for each mode of transport. MSD analysis was used to determine the minimum run length of processive runs based on the threshold used for change point analysis (Fig S3B). Processive runs with run lengths > 150 nm were further analyzed since tau did not have any apparent effect on stationary, diffusive, or processive runs < 150 nm.

#### Optical trapping

Optical trapping was performed similar to motility assays, except using a 10 W, 1064 nm laser (IPG Photonics) to trap phagosomes, a quadrant photodiode detector (QPD) (PDP90A, Thorlabs) to sense the displacement and forces exerted by phagosomes away from the trap center, and TIRF microscopy to position phagosomes over microtubules. The trap stiffness and positional calibration was determined by fitting a Lorentzian function to the power spectrum of thermal fluctuations of trapped phagosomes. Forces were calculated by measuring the displacement of the trapped phagosome from the trap center, where the displacement is directly proportional to the trap stiffness. Forces were filtered by stall time and analyzed using custom MATLAB codes to measure the magnitudes, frequency of events, and the relative binding and unbinding rates (Berger et al., 2019; Chaudhary et al., 2019).

### Quantification and statistical analysis

#### Estimation of the number of motors on cargo

To estimate the number and ratio of motors on individual cargo, we first calculated the fraction of beads that were enveloped in a phagosome (*F*_{*Ph*+}_). For each image, beads (*B*_{*TOTAL*}_) that were isolated and adequately distanced from neighboring beads so to ensure that fluorescence signals did not overlap were counted. Beads that were immunolabeled for dynein, kinesin-1, kinesin-2 or any combination of these motors were considered phagosomes (*B*_{+}_). The fraction of beads that were phagosomes was determined as follows:

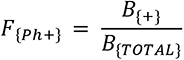

We then corrected the number of beads that were immunolabeled to determine the number of phagosomes that were positive for a given motor, where Ph*_M_* is the number of phagosomes with a positive signal for *M* motor (Fig S1H). Bootstrapping was used to determine the mean and 95% confidence intervals of the total number of phagosomes bound by each motor *M*.

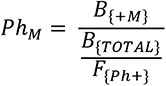

To determine the mean number of each motor (*M*) on individual phagosomes, we first calculated the number of phagosomes that were bound by one or more motors of a given type as follows:

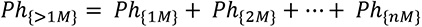

Where *Ph*_{1*M*}_ is the number of phagosomes that contain 1 *M* motor, *Ph*_{2*M*}_ is the number of phagosomes that contain 2 *M* motors, *Ph*_{*nM*}_ is the number of phagosomes that contain *n* number of *M* motors, and *Ph*_{>1*M*}_ is the total number of phagosomes that contain 1 – *n* number of *M* motors. Next, we calculated the number of phagosomes that were not bound by a given motor *M* as follows:

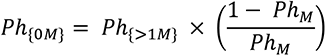

Where *Ph*_{0*M*}_ is the number of phagosomes that were not bound by a given motor *M* but bound by other types of motors. The total number of phagosomes bound by 0 – *n* number of each type of *M* motor was calculated as follows:

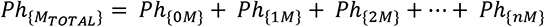

Next, we determined the total number of each motor (*M_TOTAL_*) among the total number of phagosomes (*Ph*_{*M_TOTAL_*}_) then calculated the mean number of motors (*μ*_{*M_TOTAL_*}_) on individual phagosomes using bootstrapping as follows:

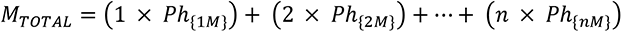

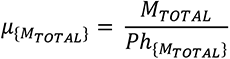

#### Estimation of the total number of motors on each cargo

To calculate the total number of motors on individual cargo, we generated a population of simulated cargoes from our experimental data. Stochastic combinations of motor type and number were generated for individual cargo using random sampling along the cumulative distribution functions of each motor (Fig S2G). Bootstrapping was used to determine the mean and 95% confidence intervals of the total number of motors on each cargo (Fig S2H).

#### Statistical analysis

All data were presented with error bars indicating standard error of the mean (SEM) or 95% confidence intervals when specified in the figure legends. All *n* and number of replicates were mentioned in the figure legends. Bootstrapping analysis was performed in MATLAB and used to test for statistical significance and determine confidence intervals. Fisher’s exact test was used to determine the significance of the frequency of plus-end and minus-end directed EPs that paused, passed, or detached from the microtubule when they encountered tau. Kolmogrov-Smirnov test was used to test for statistical significance of tau’s effect on the distribution of run lengths, velocities, force events, and relative binding rates.

## Data and code availability

All data reported and codes written in this study will be shared by the lead contact upon request.

## Acknowledgements

The authors thank Dr. Gary Brouhard for providing reagents and Dr. Florian Berger for providing assistance with unbinding data analysis. We thank Mahmoud Nour for providing assistance with the change point analysis. This work was supported by NIH.

## References

1. Akhmanova A, Hammer JA 3rd. Linking molecular motors to membrane cargo. Curr Opin Cell Biol. 2010 Aug;22(4):479–87.

2. Arimura N, Kimura T, Nakamuta S, Taya S, Funahashi Y, Hattori A, Shimada A, Ménager C, Kawabata S, Fujii K, Iwamatsu A, Segal RA, Fukuda M, Kaibuchi K. Anterograde transport of TrkB in axons is mediated by direct interaction with Slp1 and Rab27. Dev Cell. 2009 May;16(5):675–86.

3. Arpağ G, Norris SR, Mousavi SI, Soppina V, Verhey KJ, Hancock WO, Tüzel E. Motor Dynamics Underlying Cargo Transport by Pairs of Kinesin-1 and Kinesin-3 Motors. Biophys J. 2019 Mar 19;116(6):1115–1126.

4. Balabanian L, Berger CL, Hendricks AG. Acetylated Microtubules Are Preferentially Bundled Leading to Enhanced Kinesin-1 Motility. Biophys J. 2017 Oct 3;113(7):1551–1560.

5. Bentley M, Decker H, Luisi J, Banker G. A novel assay reveals preferential binding between Rabs, kinesins, and specific endosomal subpopulations. J Cell Biol. 2015 Feb 2;208(3):273–81.

6. Berger F, Klumpp S, Lipowsky R. Force-Dependent Unbinding Rate of Molecular Motors from Stationary Optical Trap Data. Nano Lett. 2019 Apr 10;19(4):2598–2602.

7. Bielska E, Schuster M, Roger Y, Berepiki A, Soanes DM, Talbot NJ, Steinberg G. Hook is an adapter that coordinates kinesin-3 and dynein cargo attachment on early endosomes. J Cell Biol. 2014 Mar 17;204(6):989–1007.

8. Blatner NR, Wilson MI, Lei C, Hong W, Murray D, Williams RL, Cho W. The structural basis of novel endosome anchoring activity of KIF16B kinesin. EMBO J. 2007 Aug 8;26(15):3709–19.

9. Blocker A, Severin FF, Burkhardt JK, Bingham JB, Yu H, Olivo JC, Schroer TA, Hyman AA, Griffiths G. Molecular requirements for bi-directional movement of phagosomes along microtubules. J Cell Biol. 1997 Apr 7;137(1):113–29.

10. Blocker A, Griffiths G, Olivo JC, Hyman AA, Severin FF. A role for microtubule dynamics in phagosome movement. J Cell Sci. 1998 Feb;111 (Pt 3):303–12.

11. Brown CL, Maier KC, Stauber T, Ginkel LM, Wordeman L, Vernos I, Schroer TA. Kinesin-2 is a motor for late endosomes and lysosomes. Traffic. 2005 Dec;6(12):1114–24.

12. Budaitis BG, Jariwala S, Rao L, Yue Y, Sept D, Verhey KJ, Gennerich A. Pathogenic mutations in the kinesin-3 motor KIF1A diminish force generation and movement through allosteric mechanisms. J Cell Biol. 2021 Apr 5;220(4):e202004227.

13. Cantalupo G, Alifano P, Roberti V, Bruni CB, Bucci C. Rab-interacting lysosomal protein (RILP): the Rab7 effector required for transport to lysosomes. EMBO J. 2001 Feb 15;20(4):683–93.

14. Chaudhary AR, Berger F, Berger CL, Hendricks AG. Tau directs intracellular trafficking by regulating the forces exerted by kinesin and dynein teams. Traffic. 2018 Feb;19(2):111–121.

15. Chaudhary AR, Lu H, Krementsova EB, Bookwalter CS, Trybus KM, Hendricks AG. MAP7 regulates organelle transport by recruiting kinesin-1 to microtubules. J Biol Chem. 2019 Jun 28;294(26):10160–10171.

16. Chen Y, Deffenbaugh NC, Anderson CT, Hancock WO. Molecular counting by photobleaching in protein complexes with many subunits: best practices and application to the cellulose synthesis complex. Mol Biol Cell. 2014 Nov 5;25(22):3630–42.

17. Christensen JR, Kendrick AA, Truong JB, Aguilar-Maldonado A, Adani V, Dzieciatkowska M, Reck-Peterson SL. Cytoplasmic dynein-1 cargo diversity is mediated by the combinatorial assembly of FTS-Hook-FHIP complexes. Elife. 2021 Dec 9;10:e74538.

18. Desjardins M, Huber LA, Parton RG, Griffiths G. Biogenesis of phagolysosomes proceeds through a sequential series of interactions with the endocytic apparatus. J Cell Biol. 1994 Mar;124(5):677–88.

19. Desjardins M, Celis JE, van Meer G, Dieplinger H, Jahraus A, Griffiths G, Huber LA. Molecular characterization of phagosomes. J Biol Chem. 1994 Dec 23;269(51):32194–200.

20. Desjardins M, Nzala NN, Corsini R, Rondeau C. Maturation of phagosomes is accompanied by changes in their fusion properties and size-selective acquisition of solute materials from endosomes. J Cell Sci. 1997 Sep;110 (Pt 18):2303–14.

21. DeWitt MA, Chang AY, Combs PA, Yildiz A. Cytoplasmic dynein moves through uncoordinated stepping of the AAA+ ring domains. Science. 2012 Jan 13;335(6065):221–5.

22. Dixit R, Ross JL, Goldman YE, Holzbaur EL. Differential regulation of dynein and kinesin motor proteins by tau. Science. 2008 Feb 22;319(5866):1086–9.

23. Dixit R, Ross JL. Studying plus-end tracking at single molecule resolution using TIRF microscopy. Methods Cell Biol. 2010;95:543–54.

24. Feng Q, Mickolajczyk KJ, Chen GY, Hancock WO. Motor Reattachment Kinetics Play a Dominant Role in Multimotor-Driven Cargo Transport. Biophys J. 2018 Jan 23;114(2):400–409.

25. Fenton AR, Jongens TA, Holzbaur ELF. Mitochondrial adaptor TRAK2 activates and functionally links opposing kinesin and dynein motors. Nat Commun. 2021 Jul 28;12(1):4578.

26. Ferro LS, Fang Q, Eshun-Wilson L, Fernandes J, Jack A, Farrell DP, Golcuk M, Huijben T, Costa K, Gur M, DiMaio F, Nogales E, Yildiz A. Structural and functional insight into regulation of kinesin-1 by microtubule-associated protein MAP7. Science. 2022 Jan 21;375(6578):326–331.

27. Flores-Rodriguez N, Rogers SS, Kenwright DA, Waigh TA, Woodman PG, Allan VJ. Roles of dynein and dynactin in early endosome dynamics revealed using automated tracking and global analysis. PLoS One. 2011;6(9):e24479.

28. Fu MM, Holzbaur EL. JIP1 regulates the directionality of APP axonal transport by coordinating kinesin and dynein motors. J Cell Biol. 2013 Aug 5;202(3):495–508.

29. Garin J, Diez R, Kieffer S, Dermine JF, Duclos S, Gagnon E, Sadoul R, Rondeau C, Desjardins M. The phagosome proteome: insight into phagosome functions. J Cell Biol. 2001 Jan 8;152(1):165–80.

30. Goyette G, Boulais J, Carruthers NJ, Landry CR, Jutras I, Duclos S, Dermine JF, Michnick SW, LaBoissière S, Lajoie G, Barreiro L, Thibault P, Desjardins M. Proteomic characterization of phagosomal membrane microdomains during phagolysosome biogenesis and evolution. Mol Cell Proteomics. 2012 Nov;11(11):1365–77.

31. Gross SP, Welte MA, Block SM, Wieschaus EF. Coordination of opposite-polarity microtubule motors. J Cell Biol. 2002 Feb 18;156(4):715–24.

32. Guardia CM, Farías GG, Jia R, Pu J, Bonifacino JS. BORC Functions Upstream of Kinesins 1 and 3 to Coordinate Regional Movement of Lysosomes along Different Microtubule Tracks. Cell Rep. 2016 Nov 15;17(8):1950–1961.

33. Gumy LF, Katrukha EA, Grigoriev I, Jaarsma D, Kapitein LC, Akhmanova A, Hoogenraad CC. MAP2 Defines a Pre-axonal Filtering Zone to Regulate KIF1-versus KIF5-Dependent Cargo Transport in Sensory Neurons. Neuron. 2017 Apr 19;94(2):347–362.e7.

34. Gyparaki MT, Arab A, Sorokina EM, Santiago-Ruiz AN, Bohrer CH, Xiao J, Lakadamyali M. Tau forms oligomeric complexes on microtubules that are distinct from tau aggregates. Proc Natl Acad Sci U S A. 2021 May 11;118(19):e2021461118.

35. Harterink M, Edwards SL, de Haan B, Yau KW, van den Heuvel S, Kapitein LC, Miller KG, Hoogenraad CC. Local microtubule organization promotes cargo transport in C. elegans dendrites. J Cell Sci. 2018 Oct 22;131(20):jcs223107.

36. Hendricks AG, Goldman YE, Holzbaur EL. Reconstituting the motility of isolated intracellular cargoes. Methods Enzymol. 2014;540:249–62.

37. Hendricks AG, Perlson E, Ross JL, Schroeder HW 3rd, Tokito M, Holzbaur EL. Motor coordination via a tug-of-war mechanism drives bidirectional vesicle transport. Curr Biol. 2010 Apr 27;20(8):697–702.

38. Hendricks AG, Holzbaur EL, Goldman YE. Force measurements on cargoes in living cells reveal collective dynamics of microtubule motors. Proc Natl Acad Sci U S A. 2012 Nov 6;109(45):18447–52.

39. Henrichs V, Grycova L, Barinka C, Nahacka Z, Neuzil J, Diez S, Rohlena J, Braun M, Lansky Z. Mitochondria-adaptor TRAK1 promotes kinesin-1 driven transport in crowded environments. Nat Commun. 2020 Jun 19;11(1):3123.

40. Hinrichs MH, Jalal A, Brenner B, Mandelkow E, Kumar S, Scholz T. Tau protein diffuses along the microtubule lattice. J Biol Chem. 2012 Nov 9;287(46):38559–68.

41. Hirokawa N, Noda Y, Tanaka Y, Niwa S. Kinesin superfamily motor proteins and intracellular transport. Nat Rev Mol Cell Biol. 2009 Oct;10(10):682–96.

42. Hoepfner S, Severin F, Cabezas A, Habermann B, Runge A, Gillooly D, Stenmark H, Zerial M. Modulation of receptor recycling and degradation by the endosomal kinesin KIF16B. Cell. 2005 May 6;121(3):437–50.

43. Hoeprich GJ, Thompson AR, McVicker DP, Hancock WO, Berger CL. Kinesin’s neck-linker determines its ability to navigate obstacles on the microtubule surface. Biophys J. 2014 Apr 15;106(8):1691–700.

44. Hoeprich GJ, Mickolajczyk KJ, Nelson SR, Hancock WO, Berger CL. The axonal transport motor kinesin-2 navigates microtubule obstacles via protofilament switching. Traffic. 2017 May;18(5):304–314.

45. Hooikaas PJ, Martin M, Mühlethaler T, Kuijntjes GJ, Peeters CAE, Katrukha EA, Ferrari L, Stucchi R, Verhagen DGF, van Riel WE, Grigoriev I, Altelaar AFM, Hoogenraad CC, Rüdiger SGD, Steinmetz MO, Kapitein LC, Akhmanova A. MAP7 family proteins regulate kinesin-1 recruitment and activation. J Cell Biol. 2019 Apr 1;218(4):1298–1318.

46. Horgan CP, Hanscom SR, Jolly RS, Futter CE, McCaffrey MW. Rab11-FIP3 links the Rab11 GTPase and cytoplasmic dynein to mediate transport to the endosomal-recycling compartment. J Cell Sci. 2010 Jan 15;123(Pt 2):181–91.

47. Hummel JJA, Hoogenraad CC. Inducible manipulation of motor-cargo interaction using engineered kinesin motors. J Cell Sci. 2021 Aug 1;134(15):jcs258776.

48. Hyman A, Drechsel D, Kellogg D, Salser S, Sawin K, Steffen P, Wordeman L, Mitchison T. Preparation of modified tubulins. Methods Enzymol. 1991;196:478–85.

49. Jenkins B, Decker H, Bentley M, Luisi J, Banker G. A novel split kinesin assay identifies motor proteins that interact with distinct vesicle populations. J Cell Biol. 2012 Aug 20;198(4):749–61.

50. Jordens I, Fernandez-Borja M, Marsman M, Dusseljee S, Janssen L, Calafat J, Janssen H, Wubbolts R, Neefjes J. The Rab7 effector protein RILP controls lysosomal transport by inducing the recruitment of dynein-dynactin motors. Curr Biol. 2001 Oct 30;11(21):1680–5.

51. Kellogg EH, Howes S, Ti SC, Ramírez-Aportela E, Kapoor TM, Chacón P, Nogales E. Near-atomic cryo-EM structure of PRC1 bound to the microtubule. Proc Natl Acad Sci U S A. 2016 Aug 23;113(34):9430–9.

52. Kendrick AA, Dickey AM, Redwine WB, Tran PT, Vaites LP, Dzieciatkowska M, Harper JW, Reck-Peterson SL. Hook3 is a scaffold for the opposite-polarity microtubule-based motors cytoplasmic dynein-1 and KIF1C. J Cell Biol. 2019 Sep 2;218(9):2982–3001.

53. Lipka J, Kapitein LC, Jaworski J, Hoogenraad CC. Microtubule-binding protein doublecortin-like kinase 1 (DCLK1) guides kinesin-3-mediated cargo transport to dendrites. EMBO J. 2016 Feb 1;35(3):302–18.

54. Liu JS, Schubert CR, Fu X, Fourniol FJ, Jaiswal JK, Houdusse A, Stultz CM, Moores CA, Walsh CA. Molecular basis for specific regulation of neuronal kinesin-3 motors by doublecortin family proteins. Mol Cell. 2012 Sep 14;47(5):707–21.

55. Loubéry S, Wilhelm C, Hurbain I, Neveu S, Louvard D, Coudrier E. Different microtubule motors move early and late endocytic compartments. Traffic. 2008 Apr;9(4):492–509.

56. Mandelkow EM, Stamer K, Vogel R, Thies E, Mandelkow E. Clogging of axons by tau, inhibition of axonal traffic and starvation of synapses. Neurobiol Aging. 2003 Dec;24(8):1079–85.

57. Matsushita M, Tanaka S, Nakamura N, Inoue H, Kanazawa H. A novel kinesin-like protein, KIF1Bbeta3 is involved in the movement of lysosomes to the cell periphery in non-neuronal cells. Traffic. 2004 Mar;5(3):140–51.

58. McVicker DP, Chrin LR, Berger CL. The nucleotide-binding state of microtubules modulates kinesin processivity and the ability of Tau to inhibit kinesin-mediated transport. J Biol Chem. 2011 Dec 16;286(50):42873–80.

59. McVicker DP, Hoeprich GJ, Thompson AR, Berger CL. Tau interconverts between diffusive and stable populations on the microtubule surface in an isoform and lattice specific manner. Cytoskeleton (Hoboken). 2014 Mar;71(3):184–94.

60. Monroy BY, Sawyer DL, Ackermann BE, Borden MM, Tan TC, Ori-McKenney KM. Competition between microtubule-associated proteins directs motor transport. Nat Commun. 2018 Apr 16;9(1):1487.

61. Monroy BY, Tan TC, Oclaman JM, Han JS, Simó S, Niwa S, Nowakowski DW, McKenney RJ, Ori-McKenney KM. A Combinatorial MAP Code Dictates Polarized Microtubule Transport. Dev Cell. 2020 Apr 6;53(1):60–72.e4.

62. Müller MJ, Klumpp S, Lipowsky R. Tug-of-war as a cooperative mechanism for bidirectional cargo transport by molecular motors. Proc Natl Acad Sci U S A. 2008 Mar 25;105(12):4609–14.

63. Niwa S, Tanaka Y, Hirokawa N. KIF1Bbeta- and KIF1A-mediated axonal transport of presynaptic regulator Rab3 occurs in a GTP-dependent manner through DENN/MADD. Nat Cell Biol. 2008 Nov;10(11):1269–79.

64. Ohashi KG, Han L, Mentley B, Wang J, Fricks J, Hancock WO. Load-dependent detachment kinetics plays a key role in bidirectional cargo transport by kinesin and dynein. Traffic. 2019 Apr;20(4):284–294.

65. Pu J, Guardia CM, Keren-Kaplan T, Bonifacino JS. Mechanisms and functions of lysosome positioning. J Cell Sci. 2016 Dec 1;129(23):4329–4339.

66. Rai AK, Rai A, Ramaiya AJ, Jha R, Mallik R. Molecular adaptations allow dynein to generate large collective forces inside cells. Cell. 2013 Jan 17;152(1-2):172–82.

67. Rai A, Pathak D, Thakur S, Singh S, Dubey AK, Mallik R. Dynein Clusters into Lipid Microdomains on Phagosomes to Drive Rapid Transport toward Lysosomes. Cell. 2016 Feb 11;164(4):722–34.

68. Reck-Peterson SL, Redwine WB, Vale RD, Carter AP. The cytoplasmic dynein transport machinery and its many cargoes. Nat Rev Mol Cell Biol. 2018 Jun;19(6):382–398.

69. Rosa-Ferreira C, Munro S. Arl8 and SKIP act together to link lysosomes to kinesin-1. Dev Cell. 2011 Dec 13;21(6):1171–8.

70. Ruhnow F, Zwicker D, Diez S. Tracking single particles and elongated filaments with nanometer precision. Biophys J. 2011 Jun 8;100(11):2820–8.

71. Schonteich E, Wilson GM, Burden J, Hopkins CR, Anderson K, Goldenring JR, Prekeris R. The Rip11/Rab11-FIP5 and kinesin II complex regulates endocytic protein recycling. J Cell Sci. 2008 Nov 15;121(Pt 22):3824–33.

72. Schroeder HW 3rd, Hendricks AG, Ikeda K, Shuman H, Rodionov V, Ikebe M, Goldman YE, Holzbaur EL. Force-dependent detachment of kinesin-2 biases track switching at cytoskeletal filament intersections. Biophys J. 2012 Jul 3;103(1):48–58.

73. Schuster M, Lipowsky R, Assmann MA, Lenz P, Steinberg G. Transient binding of dynein controls bidirectional long-range motility of early endosomes. Proc Natl Acad Sci U S A. 2011 Mar 1;108(9):3618–23.

74. Siahaan V, Krattenmacher J, Hyman AA, Diez S, Hernández-Vega A, Lansky Z, Braun M. Kinetically distinct phases of tau on microtubules regulate kinesin motors and severing enzymes. Nat Cell Biol. 2019 Sep;21(9):1086–1092.

75. Shastry S, Hancock WO. Neck linker length determines the degree of processivity in kinesin-1 and kinesin-2 motors. Curr Biol. 2010 May 25;20(10):939–43.

76. Soppina V, Rai AK, Ramaiya AJ, Barak P, Mallik R. Tug-of-war between dissimilar teams of microtubule motors regulates transport and fission of endosomes. Proc Natl Acad Sci U S A. 2009 Nov 17;106(46):19381–6.

77. Tan R, Lam AJ, Tan T, Han J, Nowakowski DW, Vershinin M, Simó S, Ori-McKenney KM, McKenney RJ. Microtubules gate tau condensation to spatially regulate microtubule functions. Nat Cell Biol. 2019 Sep;21(9):1078–1085.

78. Tinevez JY, Perry N, Schindelin J, Hoopes GM, Reynolds GD, Laplantine E, Bednarek SY, Shorte SL, Eliceiri KW. TrackMate: An open and extensible platform for single-particle tracking. Methods. 2017 Feb 15;115:80–90.

79. Ueno H, Huang X, Tanaka Y, Hirokawa N. KIF16B/Rab14 molecular motor complex is critical for early embryonic development by transporting FGF receptor. Dev Cell. 2011 Jan 18;20(1):60–71.

80. Vershinin M, Carter BC, Razafsky DS, King SJ, Gross SP. Multiple-motor based transport and its regulation by Tau. Proc Natl Acad Sci U S A. 2007 Jan 2;104(1):87–92.

81. Vershinin M, Xu J, Razafsky DS, King SJ, Gross SP. Tuning microtubule-based transport through filamentous MAPs: the problem of dynein. Traffic. 2008 Jun;9(6):882–92.

82. Welte MA. Bidirectional transport along microtubules. Curr Biol. 2004 Jul 13;14(13):R525–37.

83. Cella Zanacchi F, Manzo C, Magrassi R, Derr ND, Lakadamyali M. Quantifying Protein Copy Number in Super Resolution Using an Imaging-Invariant Calibration. Biophys J. 2019 Jun 4;116(11):2195–2203.

