## Supplemental Information for "The sets of kinesins and dynein transporting endocytic cargoes determine the effect of tau on their motility"

**This PDF file includes:**

Supplemental Figures S1 to S5

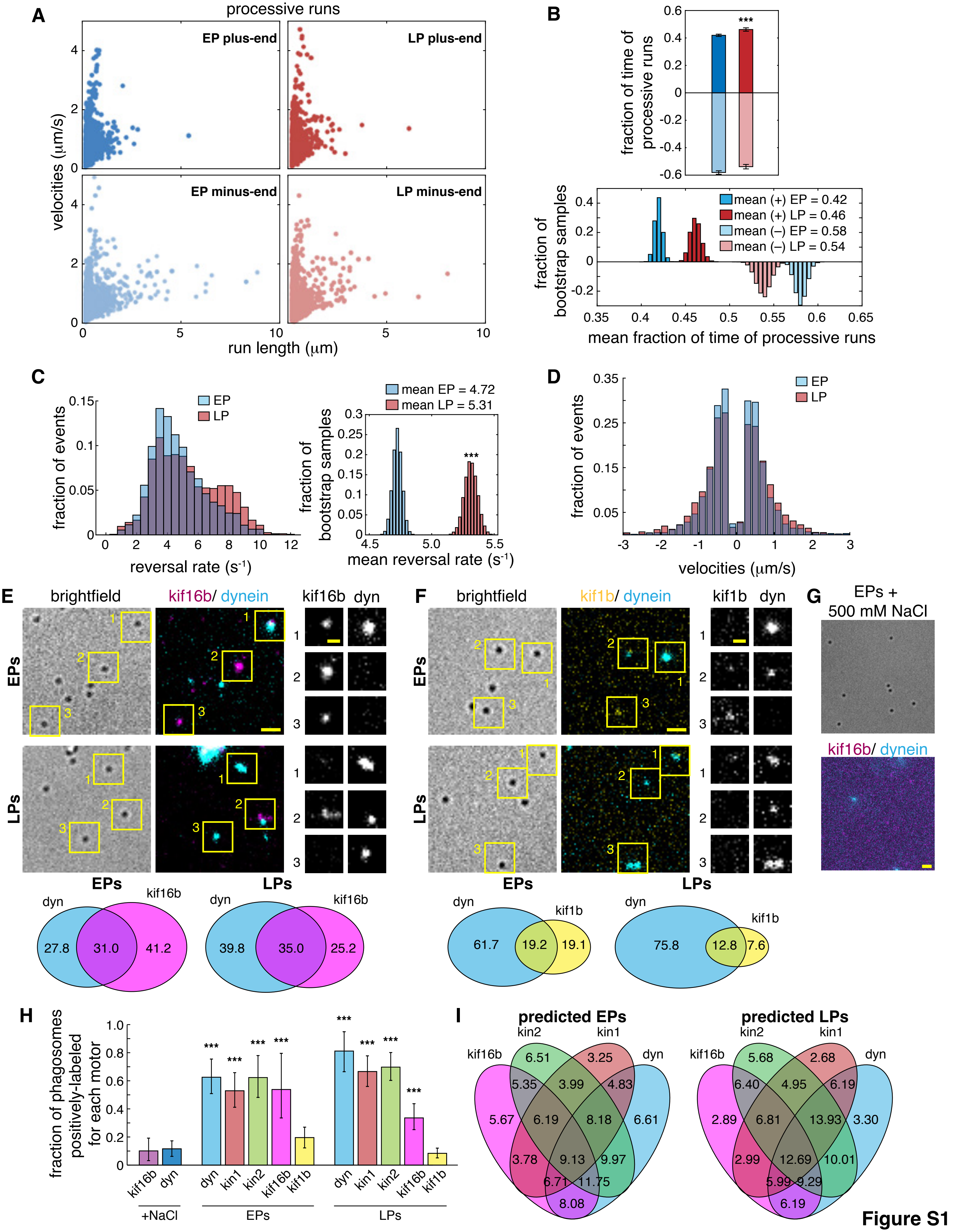
A
processive runs
EP plus-end
LP plus-end
EP minus-end
LP minus-end
velocities (μm/s)
run length (μm)
B
fraction of time of processive runs
mean (+) EP = 0.42
mean (+) LP = 0.46
mean (–) EP = 0.58
mean (–) LP = 0.54
fraction of bootstrap samples
mean fraction of time of processive runs
C
fraction of events
EP
LP
reversal rate (s-1)
mean EP = 4.72
mean LP = 5.31
fraction of bootstrap samples
mean reversal rate (s-1)
D
fraction of events
EP
LP
velocities (μm/s)
E
brightfield
kif16b/ dynein
kif16b dyn
EPs
LPs
dyn
kif16b
27.8
31.0
41.2
39.8
35.0
25.2
F
brightfield
kif1b/ dynein
kif1b dyn
EPs
LPs
dyn
kif1b
61.7
19.2
19.1
75.8
12.8
7.6
G
EPs + 500 mM NaCl
kif16b/ dynein
H
fraction of phagosomes positively-labeled for each motor
kif16b dyn
dyn kin1 kin2 kif16b kif1b
+NaCl
EPs
LPs
I
predicted EPs
predicted LPs
kif16b kin2 kin1 dyn
6.51 3.25 5.35 3.99 4.83 5.67 6.19 8.18 6.61 3.78 9.13 9.97 6.71 11.75 8.08
5.68 2.68 6.40 4.95 6.19 2.89 6.81 13.93 3.30 2.99 12.69 10.01 5.99 9.29 6.19

Figure S1

**Figure S1. Related to Figure 1.**

**A)** Plots show the velocities vs. run lengths of plus-end and minus-end directed processive runs > 150 nm for EPs (3652 plus-end runs; dark blue, 4408 minus-end runs; light blue) and LPs (2655 plus end runs; dark red, 2768 minus-end runs; light red) in cells. **B)** A bar graph shows the fraction of time of plus-end and minus-end directed processive runs for EPs and LPs. Shown below, bootstrap analysis was used to test the difference in the mean fraction of plus-end and minus-end processive motility for EPs and LPs and to determine the error bars that show 95% confidence intervals. **C)** A bar graph shows the distribution of the number of reversals for EP and LP trajectories. To the right, bootstrapping was used to determine the statistical significance between the mean reversal rates of EPs and LPs. **D)** A plot shows the velocities of processive periods of motility of EPs and LPs in the plus-end and minus-end directions. The numbers of runs are the same as in panel A. **E and F)** Images show isolated EPs and LPs immunolabeled for E) kif16b and dynein and F) kif1b and dynein. On the right, zoomed in ROIs show the different combinations of motors on individual phagosomes. Below, Venn diagrams show the mean percentages of individual motors and combinations of A) kif16b and dynein on EPs (n = 83) and LPs (n = 109) and B) kif1b and dynein on EPs (n = 103) and LPs (n = 113). Scale bars are 2 μm or 1 μm for selected ROIs. **G)** Images show phagosomes treated with 500 mM NaCl to strip proteins from the outer membranes and immunolabeled for kif16b and dynein. Salt-stripped phagosomes were used as a non-specific labeling control for immunofluorescence experiments. **H)** A table shows the corrected fraction of phagosomes immunolabeled for each motor (see methods). Bootstrapping was used to determine statistical significance (*** $p < 0.0001$). **I)** Venn diagrams show the predicted percentages of cargo bound by kinesin-1, -2, -3, dynein, or various combinations of different motors on EPs (n = 953) and LPs (n = 969). Estimates are based on random sampling of the data obtained from multi-color immunofluorescence experiments (see methods).

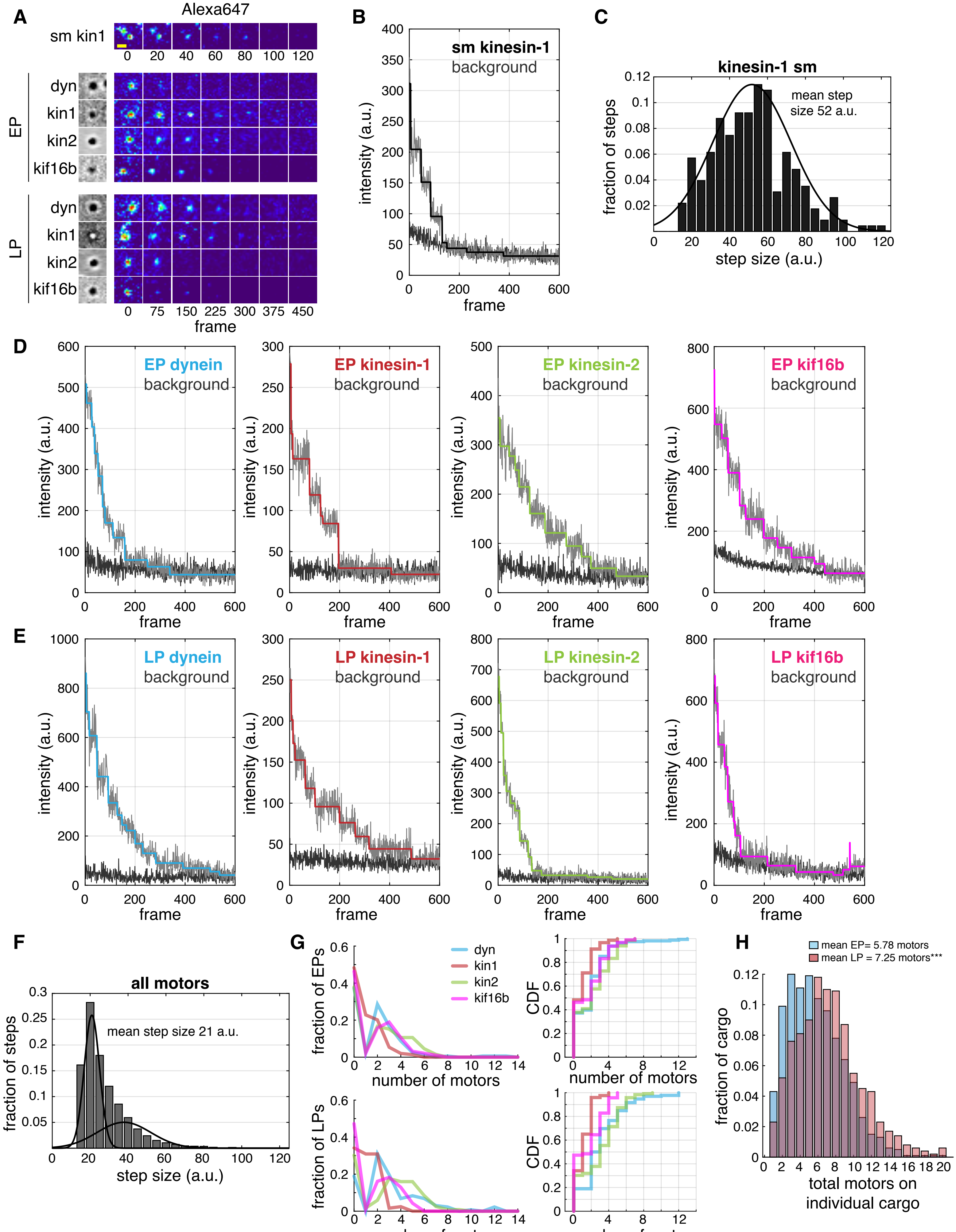

A
Alexa647
sm kin1
EP
dyn
kin1
kin2
kif16b
LP
dyn
kin1
kin2
kif16b
frame
B
sm kinesin-1
background
intensity (a.u.)
frame
C
kinesin-1 sm
mean step size 52 a.u.
fraction of steps
step size (a.u.)
D
EP dynein
EP kinesin-1
EP kinesin-2
EP kif16b
background
intensity (a.u.)
frame
E
LP dynein
LP kinesin-1
LP kinesin-2
LP kif16b
background
intensity (a.u.)
frame
F
all motors
mean step size 21 a.u.
fraction of steps
step size (a.u.)
G
dyn
kin1
kin2
kif16b
fraction of EPs
fraction of LPs
number of motors
CDF
H
mean EP= 5.78 motors
mean LP = 7.25 motors***
fraction of cargo
total motors on individual cargo


**Figure S2**

**Figure S2. Related to Figure 1.**

**A)** Time lapse images of photobleaching experiments show how the fluorescence intensity of a single kinesin-1 motor (sm kin1), or the sets of dynein (dyn), kinesin-1 (kin1), kinesin-2 (kin2), or kif16b motors on EPs and LPs decrease over time. Scale bar is 1 μm. **B)** The trace shows the stepwise decrease in fluorescence signal of a single molecule of kinesin-1. **C)** A plot shows the distribution of the step sizes for single kinesin-1 motors. The mean unitary step size was determined to be 52 a.u. **D** and **E)** Traces show the stepwise decrease in fluorescence signal compared to background for dynein, kinesin-1, kinesin-2, and kif16b on D) EPs and E) LPs. A step-finding algorithm based on the Student's t-test (Chen et al., 2014; Chaudhary et al., 2018) was used to determine the number and the size of steps for each trace. **F)** A plot shows the distribution of step sizes for kinesin-1, kinesin-2, kif16b, and dynein motors counted on EPs and LPs. A Gaussian Mixture Model was used to determine the mean step size for a single fluorophore. The mean step size was determined to be 21 a.u., which is different from the mean step size of a single motor because phagosomes were imaged using epifluorescence microscopy, whereas single motors were imaged using TRIF microscopy. **G)** Plots show the distribution of the fraction of phagosomes bound by 0–14 kinesin-1, kinesin-2, kif16b, and dynein motors. The cumulative distribution function plots are shown to the right. **H)** A histogram shows the distribution of the predicted total number of motors on individual EPs and LPs(see methods). Bootstrapping was used to calculate the means and determine the statistical significance (*** $p < 0.0001$).

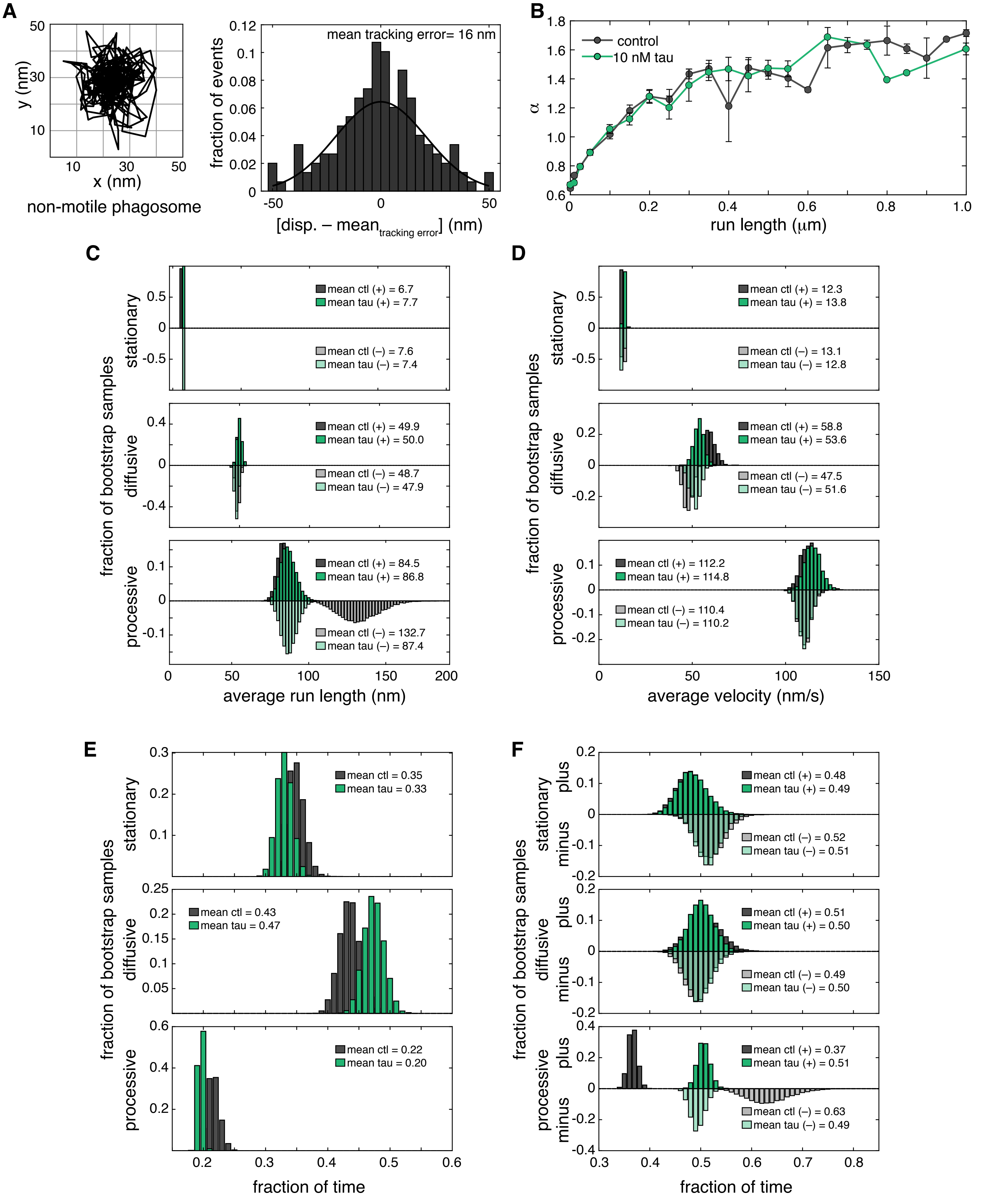

A
non-motile phagosome
x (nm)
y (nm)
mean tracking error= 16 nm
fraction of events
[disp. – mean tracking error] (nm)
B
control
10 nM tau
α
run length (μm)
C
stationary
diffusive
processive
mean ctl (+) = 6.7
mean tau (+) = 7.7
mean ctl (–) = 7.6
mean tau (–) = 7.4
mean ctl (+) = 49.9
mean tau (+) = 50.0
mean ctl (–) = 48.7
mean tau (–) = 47.9
mean ctl (+) = 84.5
mean tau (+) = 86.8
mean ctl (–) = 132.7
mean tau (–) = 87.4
fraction of bootstrap samples
average run length (nm)
D
mean ctl (+) = 12.3
mean tau (+) = 13.8
mean ctl (–) = 13.1
mean tau (–) = 12.8
mean ctl (+) = 58.8
mean tau (+) = 53.6
mean ctl (–) = 47.5
mean tau (–) = 51.6
mean ctl (+) = 112.2
mean tau (+) = 114.8
mean ctl (–) = 110.4
mean tau (–) = 110.2
average velocity (nm/s)
E
mean ctl = 0.35
mean tau = 0.33
mean ctl = 0.43
mean tau = 0.47
mean ctl = 0.22
mean tau = 0.20
fraction of time
F
plus
minus
mean ctl (+) = 0.48
mean tau (+) = 0.49
mean ctl (–) = 0.52
mean tau (–) = 0.51
mean ctl (+) = 0.51
mean tau (+) = 0.50
mean ctl (–) = 0.49
mean tau (–) = 0.50
mean ctl (+) = 0.37
mean tau (+) = 0.51
mean ctl (–) = 0.63
mean tau (–) = 0.49
fraction of time


**Figure S3**

**Figure S3. Related to Figure 3.**

**A)** A plot shows the positional tracking of a non-motile phagosome used to determine the tracking error for motility assays. On the right, a histogram shows the displacement of non-motile phagosomes (n = 9) subtracted by the mean tracking error. **B)** A plot shows the MSD of runs parsed by run length for EP +/- tau following change point analysis. The MSD was calculated to validate the change point analysis threshold used to identify diffusive and processive periods of motility. Typically, runs with lengths > 150 nm were processive ($\alpha > 1$), while shorter runs were diffusive ($\alpha < 1$). Bootstrap analysis was performed to test the impact of tau on the average **C)** run length and **D)** velocity for stationary, diffusive and processive runs, **E)** the fraction of time of stationary, diffusive, and processive runs, and **F)** the fraction of time of plus-end and minus-end directed stationary, diffusive and processive motility.

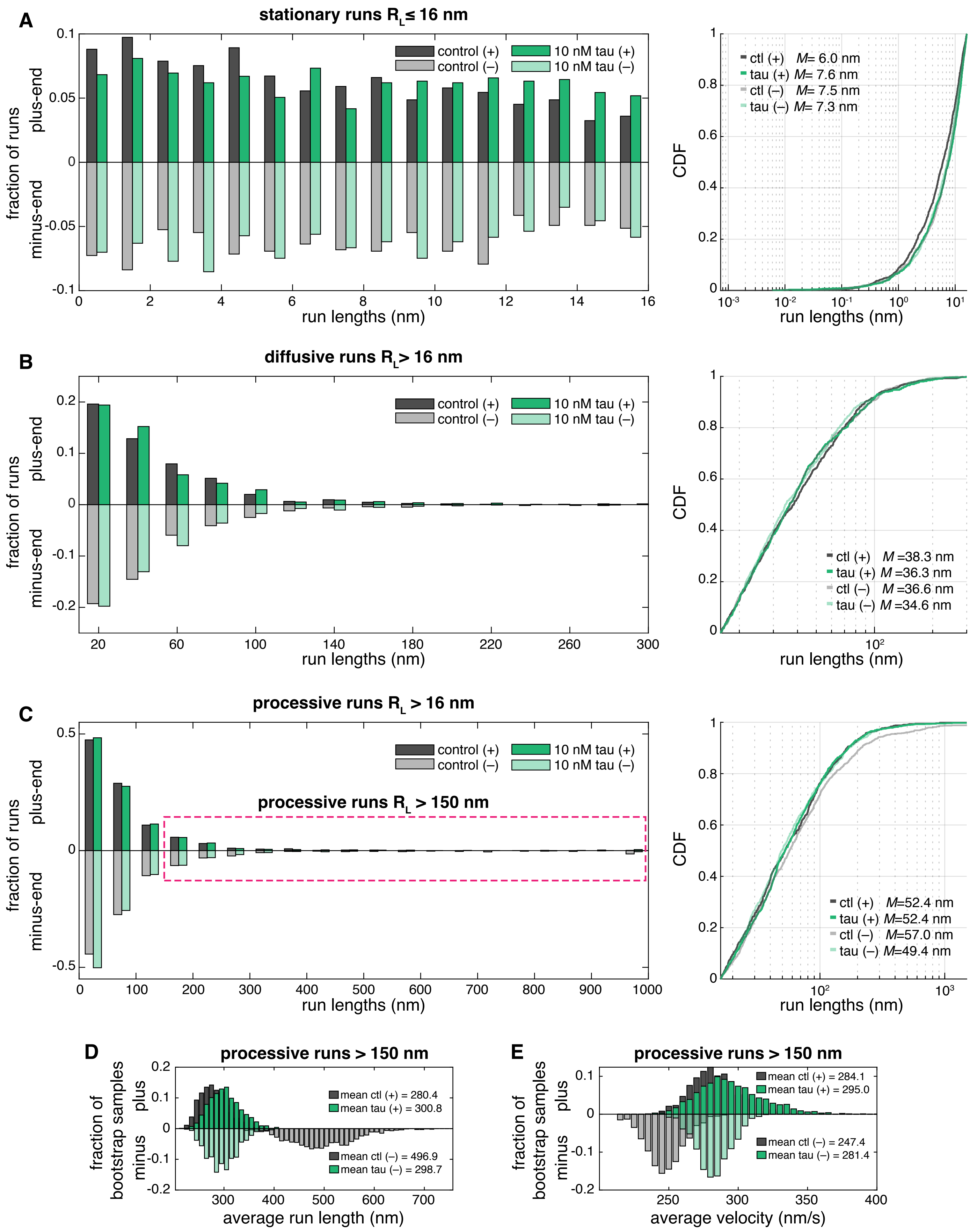
A
stationary runs R_L ≤ 16 nm
fraction of runs
plus-end
minus-end
control (+)
control (–)
10 nM tau (+)
10 nM tau (–)
run lengths (nm)
CDF
ctl (+) M= 6.0 nm
tau (+) M= 7.6 nm
ctl (–) M= 7.5 nm
tau (–) M= 7.3 nm
B
diffusive runs R_L > 16 nm
ctl (+) M =38.3 nm
tau (+) M =36.3 nm
ctl (–) M =36.6 nm
tau (–) M =34.6 nm
C
processive runs R_L > 16 nm
processive runs R_L > 150 nm
ctl (+) M=52.4 nm
tau (+) M=52.4 nm
ctl (–) M=57.0 nm
tau (–) M=49.4 nm
D
processive runs > 150 nm
fraction of bootstrap samples
plus
minus
mean ctl (+) = 280.4
mean tau (+) = 300.8
mean ctl (–) = 496.9
mean tau (–) = 298.7
average run length (nm)
E
processive runs > 150 nm
mean ctl (+) = 284.1
mean tau (+) = 295.0
mean ctl (–) = 247.4
mean tau (–) = 281.4
average velocity (nm/s)

**Figure S4**

**Figure S4. Related to Figure 4.**

**A–C)** Histograms show the distribution of minus-end and plus-end directed A) stationary, B) diffusive, and C) processive runs for EPs +/- tau identified by change point analysis. Stationary runs were identified as any run with a run length ($R_L$) $\leq$ 16 nm, which was determined by calculating the tracking error (Fig S3A). Diffusive runs were categorized as runs with $\alpha < 1$ and $R_L > 16$nm and processive runs were identified as runs with $\alpha > 1$ and $R_L > 16$nm. On the right, CDF plots show how tau impacts the frequency of stationary, diffusive, and processive run lengths. Tau reduced minus-end directed processive run lengths but did not significantly change stationary or diffusive run lengths (minus-end processive runs, $p < 0.05$). Median run lengths (*M*) are shown. Based on the MSD (Fig S3B), we analyzed the impact of tau on processive runs with lengths > 150 nm (dashed magenta box). Bootstrap analysis was performed to test tau's impact on the **D)** average run lengths and **E)** average velocities of plus-end and minus-end directed processive runs with $R_L > 150$ nm.

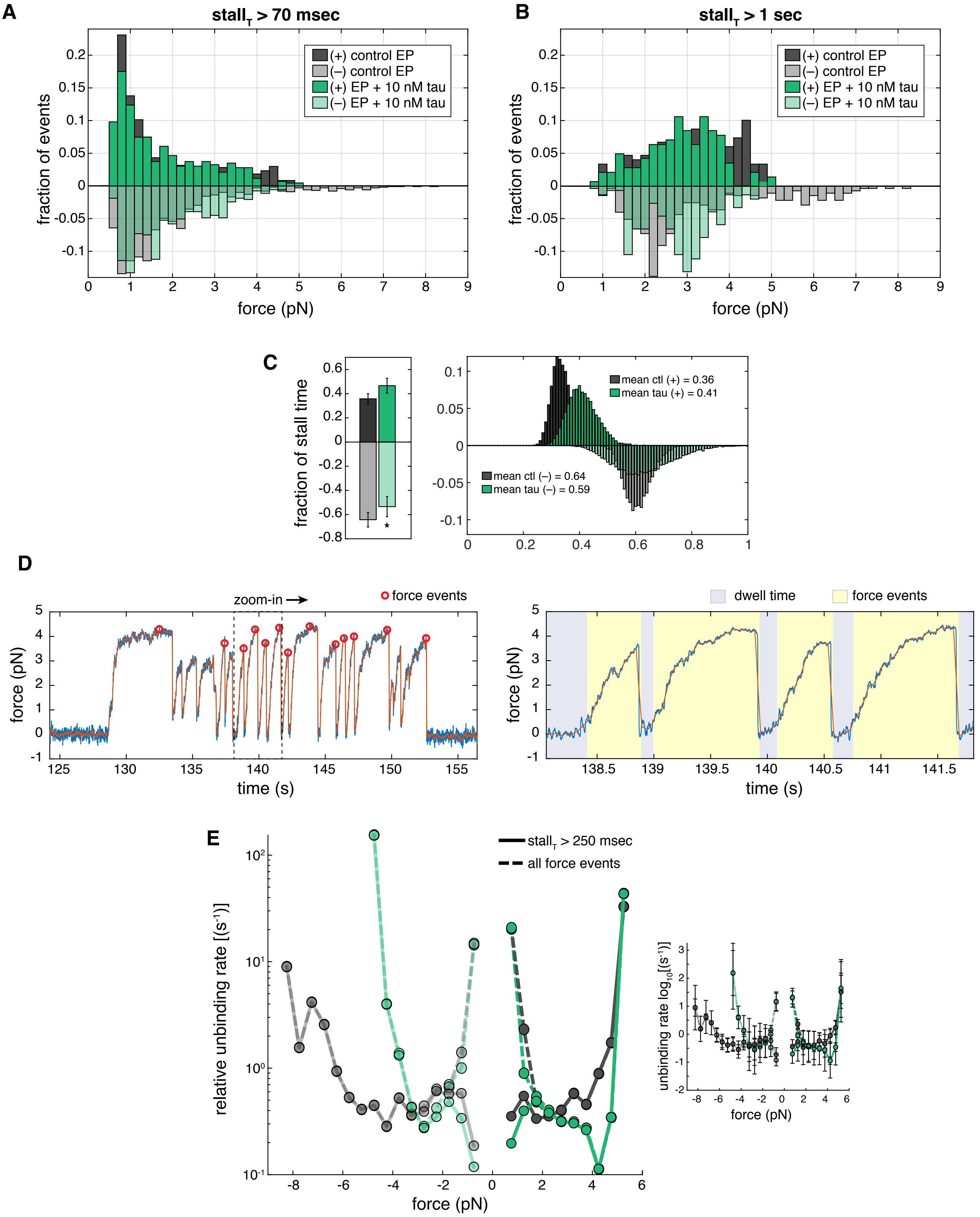
A
stall$_T$ > 70 msec
(+) control EP
(–) control EP
(+) EP + 10 nM tau
(–) EP + 10 nM tau
fraction of events
force (pN)
B
stall$_T$ > 1 sec
(+) control EP
(–) control EP
(+) EP + 10 nM tau
(–) EP + 10 nM tau
fraction of events
force (pN)
C
fraction of stall time
mean ctl (+) = 0.36
mean tau (+) = 0.41
mean ctl (–) = 0.64
mean tau (–) = 0.59
D
zoom-in
force events
force (pN)
time (s)
dwell time
force events
force (pN)
time (s)
E
stall$_T$ > 250 msec
all force events
relative unbinding rate [(s$^{-1}$)]
force (pN)
unbinding rate log$_{10}$[(s$^{-1}$)]
force (pN)

Figure S5

**Figure S5. Related to Figure 5.**

**A** and **B)** Histograms show the distribution of the total fraction of forces +/- tau in the plus-end and minus-end direction for A) events with stall times ($stall_T$) > 70 msec and B) longer sustained forces with $stall_T$ > 1 sec. Lower forces with shorter time intervals are common in both the plus-end and minus-end directions, which are thought to be due to the sets of motors detaching from the microtubule before maximum stall forces are reached. **C)** A bar graph shows the fraction of time of force events in the plus-end direction and the minus-end direction +/- tau. On the right, a plot shows how bootstrapping was used to test tau's impact on the fraction of plus-end and minus-end stall times. Means are shown (* $p < 0.05$). **D)** Plots show a force trace of a plus-end directed EP. On the left, red circles indicate force events with stall durations > 250 msec and stall forces > 0.5 pN. On the right, is a zoomed-in region from the left plot showing how the binding and unbinding rates were determined. The intervals of diffusive dwell times (purple) > 0.05 s are shown between force events (yellow). **E)** A plot shows the force-dependent relative unbinding rates calculated for plus-end and minus-end directed forces with stall durations > 250 msec (solid lines) and unfiltered forces (all force events; dashed lines) +/- tau. The plot to the right shows the log-unbinding rates on the y-axis. error bars indicate SEM.
